# Integrating remote sensing and jurisdictional observation networks to improve the resolution of ecological management

**DOI:** 10.1101/2020.06.08.140848

**Authors:** Philip A. Townsend, John Clare, Nanfeng Liu, Jennifer L. Stenglein, Christine Anhalt-Depies, Timothy R. Van Deelen, Neil A. Gilbert, Aditya Singh, Karl J. Martin, Benjamin Zuckerberg

**Author notes:** Co-lead authors (Philip A. Townsend, John Clare).

## Abstract

The emergence of citizen science, passive sensors (e.g., trail cameras and acoustic monitoring), and satellite remote sensing have enabled biological data to be collected at unprecedented spatial and temporal scales. There is growing interest in networking these datastreams to expedite the collection and synthesis of environmental and biological data to improve broad-scale ecological monitoring, but there are no examples of such networks being developed to directly inform decision-making by managing agencies. Here, we present the implementation of one such jurisdictional observation network (JON), Snapshot Wisconsin (SW), that links satellite remote sensing (RS) with a volunteer-based trail camera network to generate new insights into wildlife distributions and improve their management by the state agency. SW relies on citizen scientists to deploy trail cameras across the state and classify images of wildlife. As of early 2020 SW comprises nearly 1800 volunteers hosting >2100 active cameras recording >37 million images across a sampling effort of >2000 combined trap-years at >3300 distinct camera locations. We use a set of case studies to demonstrate the potential power of a JON to monitor wildlife with unprecedented combinations of spatial, temporal, and biological resolution and extent. Specifically, we demonstrate that SW markedly improves the spatial and temporal resolution with which black bear distributions can be monitored or forecast, in turn improving the resolution of decision-making. Enhancing the biological resolution of monitoring (e.g., monitoring the distribution of species traits or behaviors) may provide new insights into population drivers, such as the connection between vegetation productivity and white-tailed deer foraging behaviors. Enhanced taxonomic extent provided by trail cameras and other passive sensor networks provide managers new information for a wide range of species and communities that are not otherwise monitored. Our cases further show that JONs synergize existing monitoring practices by serving as a complementary and independent line of evidence or as a tool to enhance the extent and precision of existing models through integrated modeling approaches. SW and other JONS are a powerful new tool for agencies to better achieve their missions and reshape the nature of environmental decision-making.

## Introduction

Advances like the emergence of citizen science and growing use of remote sensors effective for sampling taxa locally like trail cameras, acoustic recording devices, and satellite remote sensing have enabled biological data to be collected more quickly over larger scales than ever before (Bonney et al. 2009, Steenweg et al. 2017, Shonfield and Bayne 2017). Concurrently, new satellite remote sensing (SRS) missions and openings of existing archives have unearthed a similar wealth of data related to global land cover, plant phenology, and other characteristics of Earth’s surface (Wulder et al. 2012). This combination of remotely-sensed biodiversity and environmental data at multiple scales of observation provides opportunities to improve understanding of anthropogenic impacts to biodiversity and guide decision-making.

Observation networks (ON) are programs to collect and synthesize broad-scale environmental and biological data (Keller et al. 2008, Scholes et al. 2012, Lindenmayer et al. 2018). These enterprises vary in scope, sampling structure, and data collection methodologies. Examples include eBird, which aggregates bird observations submitted by citizen scientists across the world (Sullivan et al. 2009), the urban wildlife information network, which compiles trail camera images of wildlife in cities across the US (Magle et al. 2019), or the national phenology network (Schwartz et al. 2012). The objectives of ONs are to improve ecological inference, prediction, and forecasting, and geospatial data derived from SRS play a critical operational role within ONs as a source of input variables that enable these objectives (Turner 2014). The fusion of local biodiversity observations with geospatial data provided by SRS observations is a major component of global conservation efforts like the development of essential biodiversity variables (Kissling et al. 2018) for species populations. Because these efforts strongly depend upon fusing ONs, SRS, and other geospatial datasets (Chandler et al. 2017, Jetz et al. 2019), there is substantial research interest associated with developing and implementing such integrated monitoring.

Although conservation targets are often continentally or globally defined via convention or agreement, most natural resource management and conservation decisions are made at sub-national scales (e.g., regions, provinces, counties). Decision-making across these smaller extents still must contends with biodiversity monitoring that is imperfect along taxonomic, temporal, or spatial axes (Kissling et al. 2018, Jetz et al. 2019). For example, many species are not monitored. In addition, sampling may be limited or non-representative spatially or temporally, or data may be aggregated at coarse grains that make it difficult to understand the driving changes and implement effective management solutions (Aceves-Bueno et al. 2015, Marra et al. 2015, Artelle et al. 2018, Spies et al. 2019). Concurrently, there is growing appreciation that local biodiversity changes are often driven by broader processes that can be detected via remote sensing but cannot be directly manipulated for reasons of scale (e.g., climate). Quantifying such processes and their effects on biodiversity can both improve understanding of the system state and trajectory needed to determine whether, where, and when actions are warranted (Sultaire et al. 2016, Wilson et al. 2018, Clare et al. 2019a, Spies et al. 2019).

Interest in the development of ONs operated by or in collaboration with environmental management agencies to improve decision-making has increased recently. We call these **jurisdictional observation networks** (JONs): passive monitoring efforts (*sensu* Lindenmayer and Likens 2010) that seek to fill spatial, temporal or taxonomic information gaps across a diversity of jurisdictional extents. Sampling design, methodology, and frequency may vary considerably, ranging from infrequent but spatially comprehensive biological atlases reliant upon citizen scientists (e.g., the Wisconsin Breeding Bird Atlas every 20 years) to more intensive, regular, or continuous sampling protocols facilitated by automated sensors (e.g., North Carolina’s Candid Critters). We believe effective implementation of JONs will also require the integration of broad-scale remote sensing data—including both satellite and ground sensor networks—to support biological inference and prediction across space and time.

Here, we describe implementation of one such JON that uses satellite remote sensing and a volunteer-based trail camera network to generate new insights into wildlife distributions and their management across space and time. We present Snapshot Wisconsin as proof of concept that agencies can manage the fusion of structured biodiversity observations with earth observations to collect data at unparalleled resolution and volume, which can enable agency managers and scientists to break free of the constraints of traditional monitoring approaches. We provide case studies that demonstrate the potential of fusing remote sensing and JONs to guide decision-making across a range of taxa and spatiotemporal scales.

### Snapshot Wisconsin

Snapshot Wisconsin (SW) was initiated by the Wisconsin Department of Natural Resources (WDNR) in 2014 in partnership with the University of Wisconsin and NASA with joint goals of improving information for wildlife decision-making, broadening stakeholder engagement with natural resources management, and leveraging new technologies to better characterize spatiotemporal patterns in wildlife populations. Motivation for SW reflects common monitoring limitations. Existing monitoring practices are taxonomically limited, with a primary focus on species of conservation concern or game species. Sampling constraints often compel WDNR and other agencies to employ incomplete spatial or temporal sampling based on convenience, with data typically aggregated to spatial or temporal scales that may be poorly defined or misaligned with the biological processes of interest. Reconciling differences between the extent and resolution of sampling, biological processes, and decision-making units is challenging, and often forces decision-makers to use information that is not perfectly fit for purpose (Millspaugh et al. 2009, MacFarland and Van Deelen 2011).

SW relies on citizen scientists to deploy (“host”) trail cameras across the state and to classify images. Volunteers host a single camera placed within a US Public Land Survey System (PLSS) quarter-township (comprising a 4.8 x 4.8 km cell), which is used to delineate sampling units across the state. In certain regions, denser camera placement is enacted to support specific objectives. Volunteers are expected to maintain camera stations and upload images to a central web repository multiple times per year to maintain continuous monitoring. The SW trail camera network is in many ways analogous to SRS, as the cameras operate continuously--indeed with finer temporal granularity than most SRS missions at comparable spatial scales. SW provides nonstop sampling that informs decision-making processes that occur throughout the year (see below) and facilitates broader understanding of how wildlife respond to seasonal changes in the environment (Figure 1).

**Figure 1.**
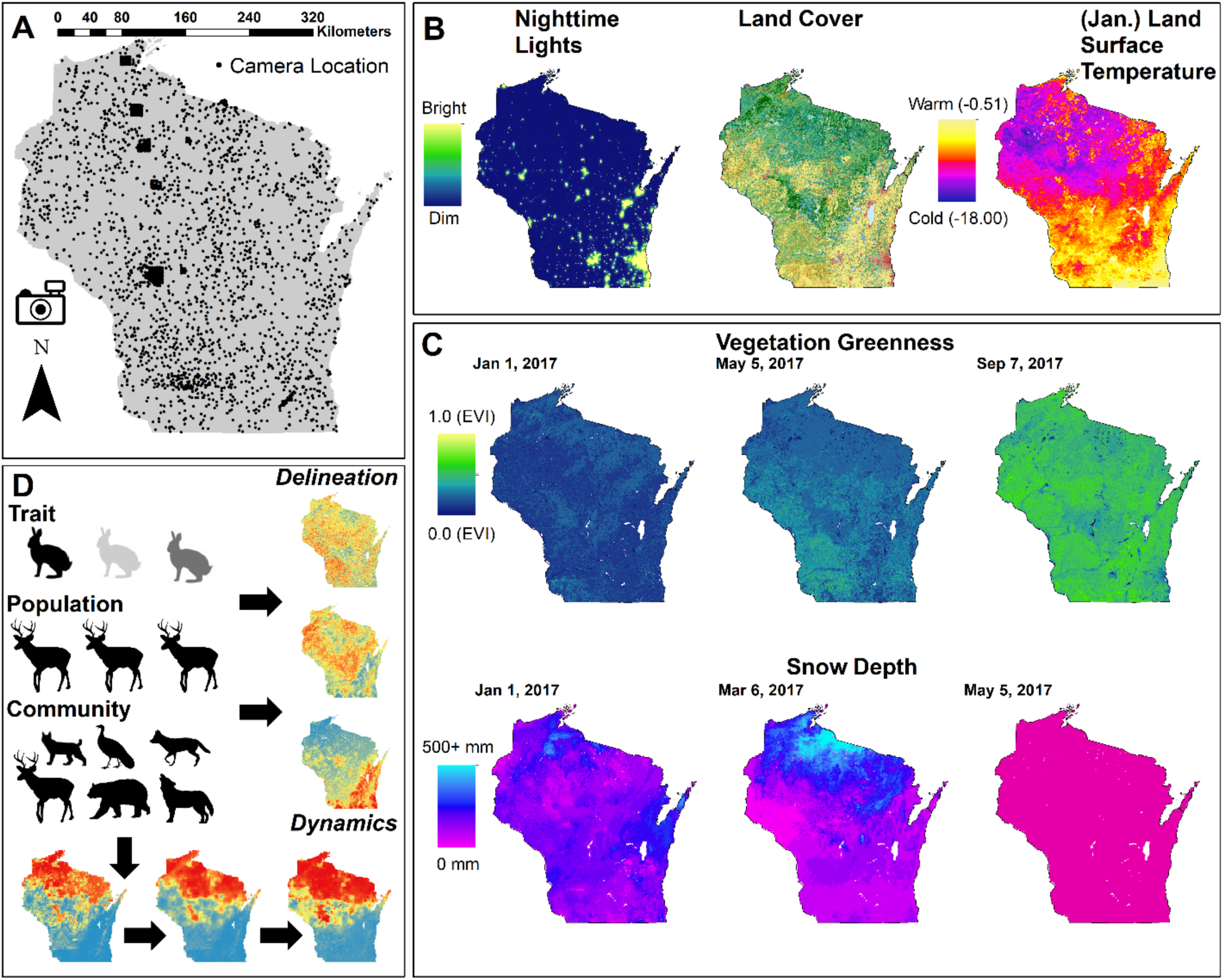
(A) The location of cameras contributing data to Snapshot Wisconsin at any point in the project’s duration through May 2020 (n = 3364). (B) Static satellite remote sensing data layers used for prediction/inference include nighttime light intensity as an indicator of human presence, Landsat-derived land cover, and longer-term averages in winter land surface temperature. (C) Dynamic spatial predictors at finer temporal resolution include vegetation greenness indices and snow depth. (D) Combining information about species traits (phenotypes, behaviors), populations, and assemblages derived from cameras with the spatial data from satellite remote sensing enables biodiversity variables to be delineated, monitored, or forecast.

SW cameras are motion-activated Bushnell Trophy Cam models (Overland Park, Kansas) and record a 3-image sequence when triggered, with a 15 s gap between triggers. Much like spaceborne remote sensing missions, SW exerts standardization to ensure data quality and compatibility across image collections. The WDNR provides cameras with fixed settings that produce encrypted photos to prevent manipulation. Furthermore, the image classification options (Appendix S1) are standardized to include: species identification, number of individuals, presence of juveniles or radio-collared animals, and behaviors exhibited in the image sequence such as foraging, vigilance, or resting. Certain sources of variation are more difficult to control via design (for example, camera placement), and instead, SW collects information about potential sources of observation variability for downstream analysis (Kelling et al. 2019). Image classification (Appendix S1) is a three-part process consisting of classifications by camera hosts, consensus-based classification by additional volunteers on the Zooniverse crowdsourcing platform (snapshotwisconsin.org), and expert evaluation that feeds into model-based strategies to account for detection and classification errors (Clare et al. 2019b), with the intent to eventually also include artificial intelligence approaches largely to remove blanks and images with humans (Willi et al. 2019).

### Integration with Satellite Remote Sensing (SRS)

Satellite remote sensing is an implicit component of SW as a basis for explaining and predicting wildlife distributions across space and time. The continuous sampling design of SW captures both intra-annual and inter-annual variability in species distributions and abundance. To test hypotheses and make predictions across temporal resolutions or extents, SW assimilates both static remote sensing products (e.g., land cover) and products that capture inter and intra-annual dynamics related to vegetation phenology and productivity. For example, temporal patterns of greenness and phenology from instruments such as MODIS can capture seasonal changes in plant productivity (and therefore food or shelter resources; Pettorelli et al. 2005). Similarly, snow cover, snow depth and frozen ground derived from remote sensing are important environmental factors explaining patterns of distribution and behavior of cold-adapted species (Zhu et al. 2019). Land cover data commonly are used to define habitat types for many species and can be used to evaluate the ecological effects of habitat loss and fragmentation (Fahrig 2003). Night lights are a surrogate for human occupancy and urbanization intensity (Mazor et al. 2013). Finally, thermal landscapes can be mapped using daytime land surface temperatures and can be effective for capturing species responses to extreme weather or longer-term climatic conditions (Albright et al. 2010). Here, we primarily focus on predictors derived from SRS to demonstrate the potential for a primarily remotely sensed approach (which is appealing because the frequency and regularity of data collection provides many benefits), but recognize that other geospatial variables may also be useful predictors (e.g., road networks).

SW has been a massive success with respect to data collection and volunteer participation. As of 2020, >37 million images have been recorded across a sampling effort of >2000 combined trap-years at more than 3300 distinct camera locations, with more than 6 million images currently known to contain wildlife (Figure 1). Nearly 1800 volunteers host >2100 active cameras, with >7,000 other volunteers assisting with image classification on the project’s crowdsourcing platform. In terms of overall sampling effort, SW is, to the best of our knowledge, the largest single continuous, self-contained camera trapping effort to date. We present the following case studies (Table 1) as examples of the power and utility of incorporating SRS data and JONs for increasing the resolution and effectiveness of biological monitoring. The SRS variables used in the work presented here are outlined in the supplementary appendix for each analysis.

**Table 1.**
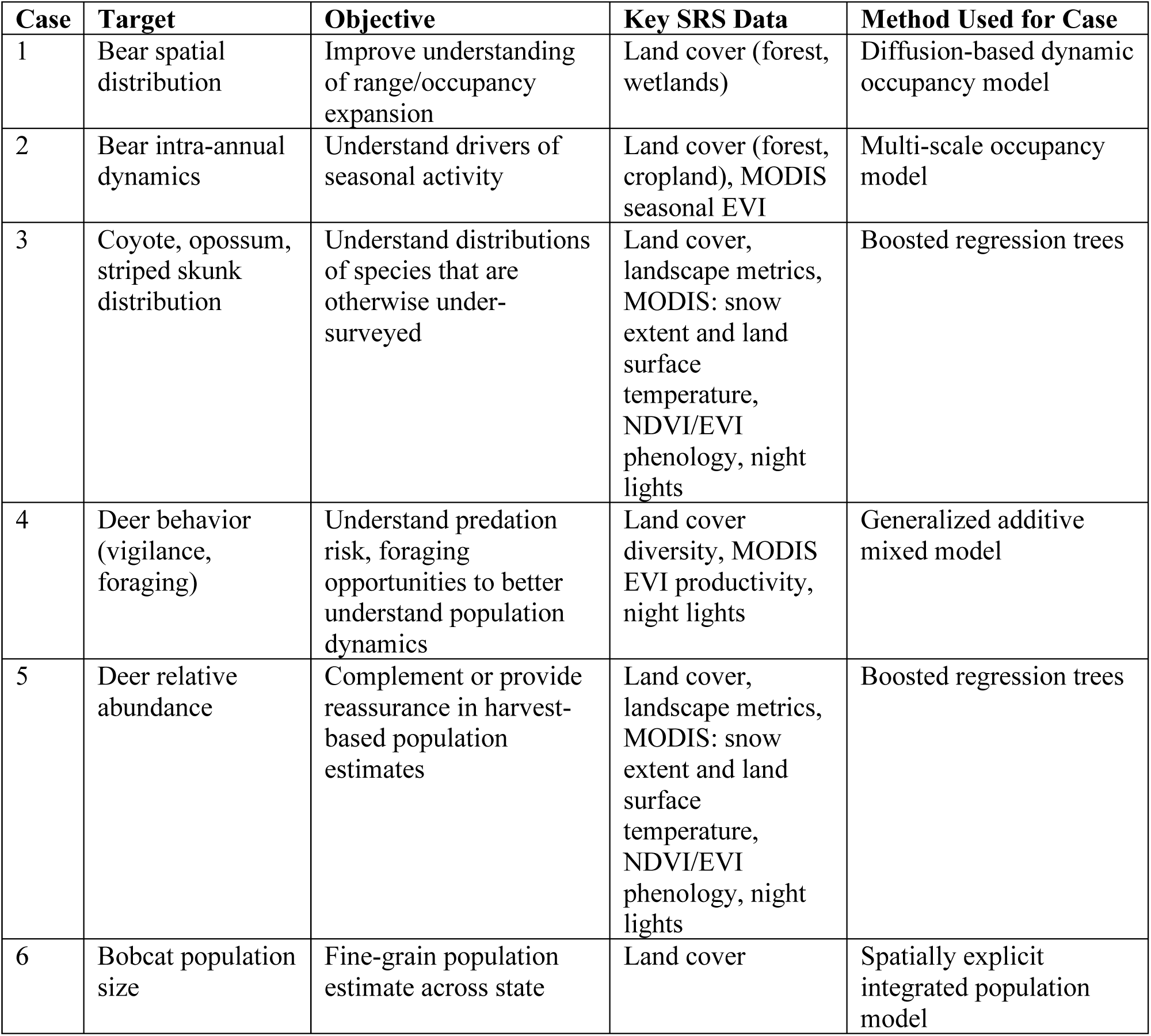
Cases presented in the study.

### Case 1. Increasing spatial resolution

Species that are of significant managerial interest are subject to more regular and rigorous monitoring by agencies, but existing monitoring programs may lack spatial or temporal resolution to answer targeted questions. For example, black bear (*Ursus americanus*) distribution is believed to be expanding into the more populated and agricultural regions of southern Wisconsin (WDNR 2019), which brings increased potential for conflict and property damage. There is interest in predicting bear population spread to anticipate conflict or potential regulatory changes, but the models to estimate population trends using animal harvest data can only characterize changes in population size at a very course grain (Allen et al. 2018a). Linking SRS data with observations from SW provides the resolution to address these needs. Predictions from a diffusion-based dynamic occupancy model at a 5 x 5 km resolution reflecting the home range size of female bears suggest a stable or potentially declining distribution from 2015-2018 (Figure 2, methods in Appendix S2), consistent with estimated population decline during the same time period using different techniques (Allen et al. 2018a). Forecasts suggest that bears are unlikely to become established within southern Wisconsin in the near future (i.e., by 2024) assuming current conditions hold, because this region is less forested, which reduces the probability of bear persistence (Figure 2, Table S2.1, Appendix S2). However, because both colonization and persistence processes appear strongly related to the occupancy state of neighboring cells, bear colonization of these regions could be rapid if occupancy reaches a critical mass.

**Figure 2.**
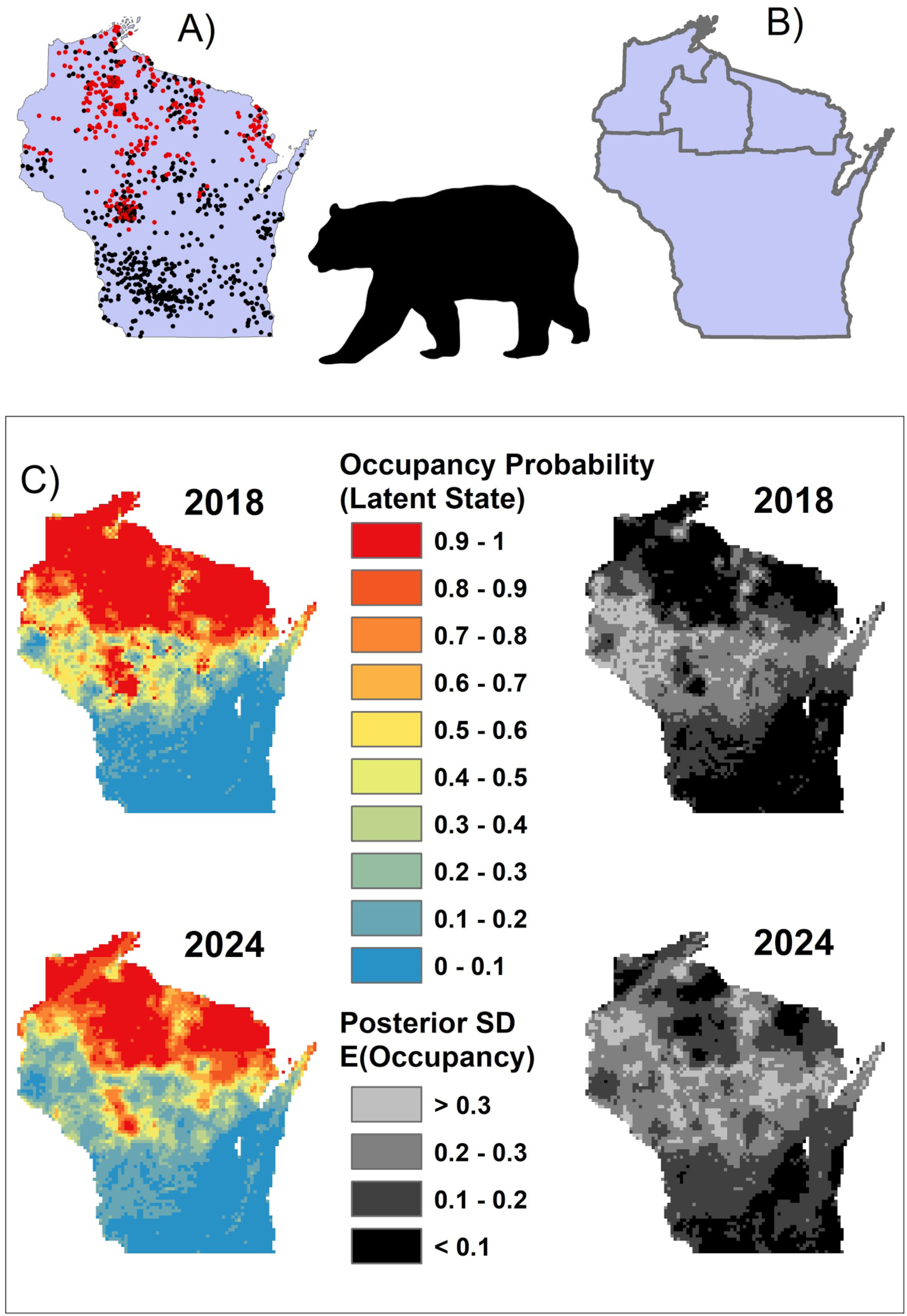
(A) Detection (red)/non-detection (black) of bears in Snapshot Wisconsin cameras from 2018; (B) coarse spatial scale bear harvest monitoring units; (C) predicted bear species occupancy and (D) associated uncertainty for 2018 and forecast for 2024, providing more detail than if summarized by management unit.

### Case 2. Increasing temporal resolution

For many mammal species, linking locally-sensed and SRS data can support decision-making at finer and potentially more useful temporal resolutions than static or annual maps. Species distributions and their environmental associations change throughout the course of the year (Conn et al. 2015, Zuckerberg et al. 2016). These dynamics are often overlooked in favor of annual maps of species distributions or long-term population trends, but management or conservation actions may be more effective and efficient when tailored to reflect changes in seasonal activity and its associated drivers. For example, deterrence actions aimed at reducing livestock depredation or crop damage are likely to be more effective when implemented in the places and seasons with a high probability of conflict (e.g., Olson et al. 2019). Seasonal patterns of black bear occurrence strongly reflect the phenology of the species and its winter torpor (Figure 3, Appendix S3). The primary drivers of bear distribution dynamics across the year include positive associations with forest cover across a broader surrounding area, and finer grained associations with concurrent daily estimates of the enhanced vegetation index (EVI, smoothed to daily values following Beck et al. 2006). Associations between bear occurrence and local forest and cropland cover exhibit some season variation: cropland appears to be avoided during peak summer months, but not during spring or autumn (Figure S3.1, Appendix S3). The likelihood of bear occurrence across the year is predicted as greatest within the northwestern part of the state, which coincides with where WDNR has made recent changes to harvest delineations to address increased conflict.

**Figure 3.**
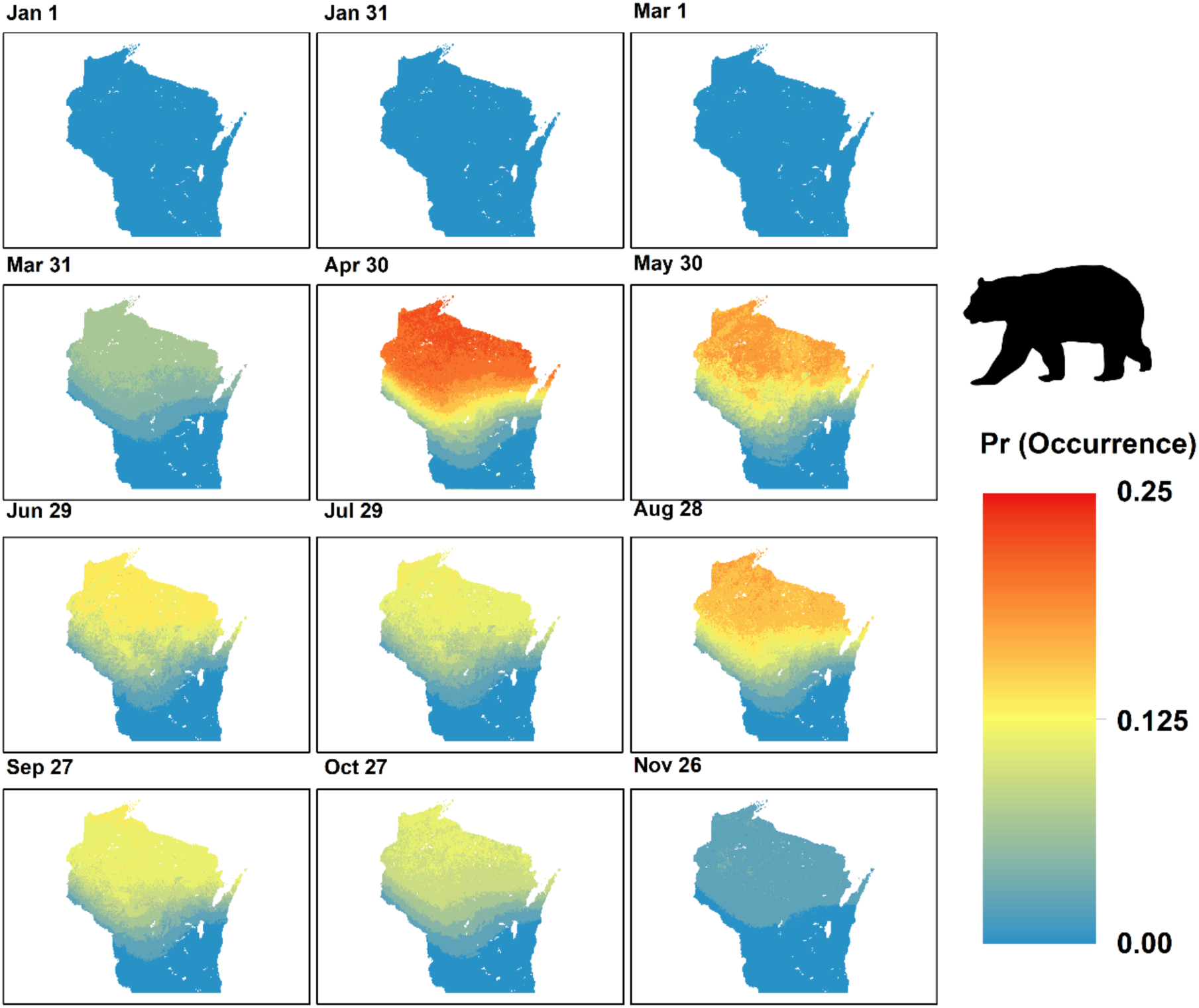
Use of continuous RS and SW data to predict bear distributions (the probability of occupancy times the daily probability of availability for detection conditional upon occupancy) at fine resolutions over the course of 2017.

### Case 3. Expanding taxonomic extent

A key contribution of SW has been the project’s capacity to provide information about species that are not otherwise monitored. Species believed to be extirpated or incidental across the state (e.g., moose, cougar, lynx) are difficult to monitor by virtue of their scarcity. Because SW’s sampling effort is extensive, continuous and quantifiable, it provides stronger evidence for scarcity than many large-scale sampling designs that operate opportunistically (Bayraktarov et al. 2019, Kays et al. 2020). For example, as of early 2020, SW has generated < 10 and 2 detections of moose and cougar, respectively, suggesting that these species of potential concern are extremely rare or incidental in Wisconsin. Similarly, SW enables the monitoring of common species that may have managerial importance aside from sporting interests or conservation concerns, but are not otherwise monitored or managed due to resource constraints and existing prioritizations. Unmonitored species such as coyote (*Canis latrans*), opossum (*Didelphis virginiana*), and striped skunk (*Mephitis mephitis*) are readily detected by SW and their distributions can be delineated and monitored at high resolution (Figure 4, Appendix S4). Each species appears to be more common in southern Wisconsin. Patterns in their occurrence appear associated with differences in land cover composition and climatic conditions between the northern and southern parts of the state, but stronger insights into the underlying drivers of each species distribution may require dynamic models (Yackulic et al. 2015). Importantly, these species may play important direct and indirect roles in the spread and prevalence of different diseases (e.g., Levi et al. 2016), and active monitoring could inform wildlife management aimed at managing human or wildlife disease risk (e.g., Rohr et al. 2020). More broadly, stronger knowledge of the distributions and habitat associations of a larger pool of species is likely to enable better quantification of biodiversity hotspots and allow biodiversity protections to be more effectively delineated (Falconer and Ford 2020). Notably, the three examples we used here are predators for important game species. Concurrent analysis of multiple species facilitates understanding wildlife community structure, diversity and species interactions (Burton et al. 2012, Allen et al. 2019).

**Figure 4.**
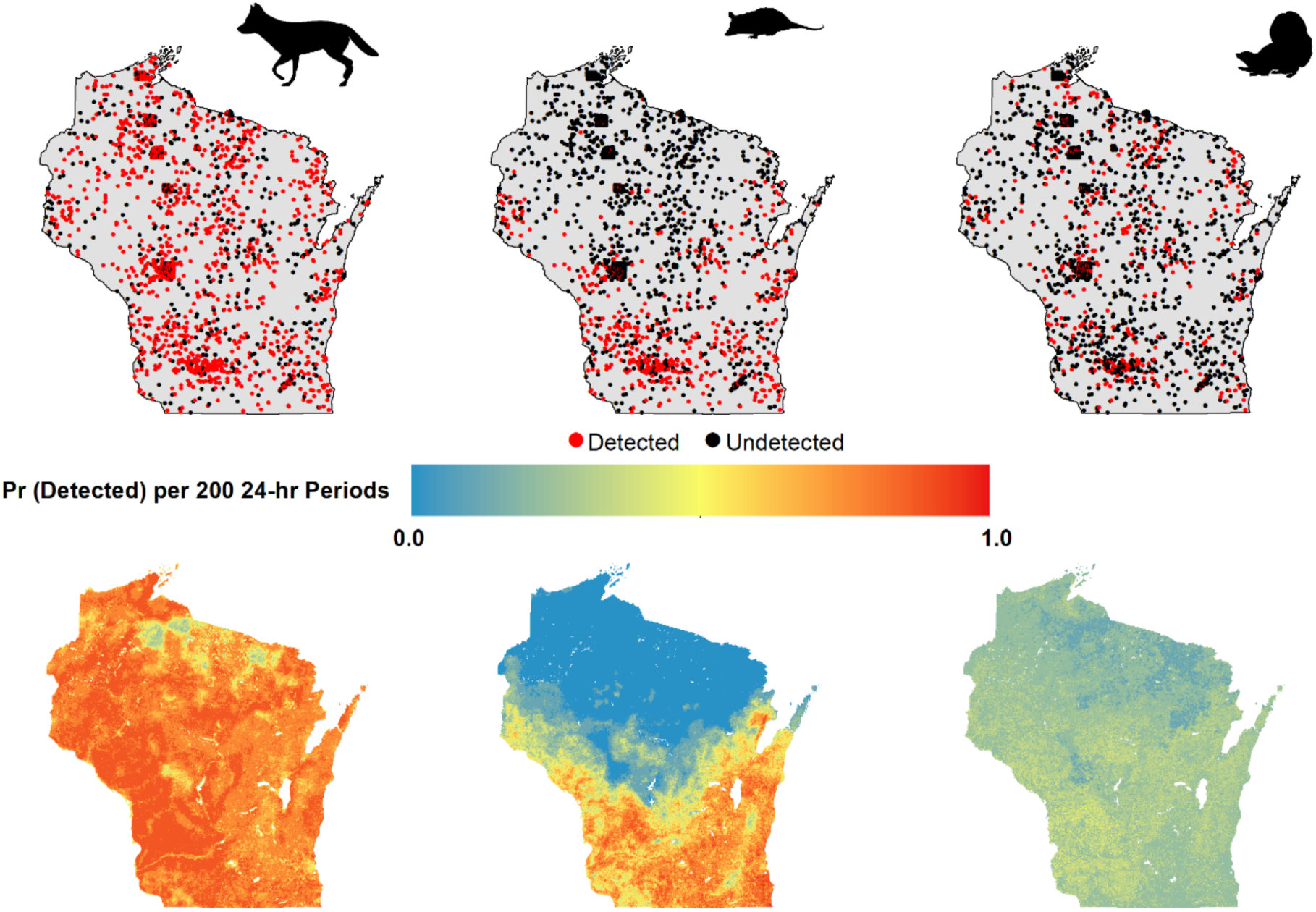
Trail camera and RS based predictions of distributions of species that are not otherwise monitored: coyote, opossum, and striped skunk. Top row: SW observations (detection/non-detection) across 2015 - 2018; bottom row: predicted distributions.

### Case 4. Increased biological resolution

SW supports decision-making and ecological understanding by providing information at increased biological resolution beyond species detection/non-detection. Trail camera images contain information about species traits or behaviors that provide insights into the mechanisms underlying population changes (e.g., Zimova et al. 2020). For example, the complexity of deer population dynamics in Wisconsin poses a challenge for effective management, with both bottom-up (food) and top-down (natural or anthropogenic predation) drivers believed to have important effects (e.g., Warbington et al. 2015). Top-down effects may further manifest in varying ways: for example, predators may influence prey populations directly via consumption or indirectly by intimidating prey and reducing their foraging efficiency (Brown and Kotler 2004, Creel 2018). Consequently, understanding the behavioral landscape and how it is shaped by predation risk and foraging opportunities can provide insights into underlying population variation and can help guide management of the system (e.g., the efficacy of predator control). SRS can play a critical role in delineating spatial and temporal variation in forage resources. For example, an analysis of deer foraging and vigilance behavior (ignoring any proxies for predation risk) suggests that deer spend relatively less time foraging in areas with greater land cover diversity and higher plant productivity (using annually-integrated MODIS EVI as a proxy; Appendix S5). Conversely, deer in predator-rich northern Wisconsin spend relatively less time being vigilant and more time foraging (Figure 5). The predicted spatial pattern suggests that deer in northern Wisconsin operate under greater nutritional stress and may suggest that deer perceive resource limitation as a greater risk than predation, although further research directly considering potential metrics of risk is needed. Moreover, the predicted incidence of foraging exhibits intra-annual variation and peaks during early spring when nutritional demands are also peaking and in late summer when mature fawns venture more broadly. This suggests that any potential non-consumptive predator effects might be more important during these specific times of year.

**Figure 5.**
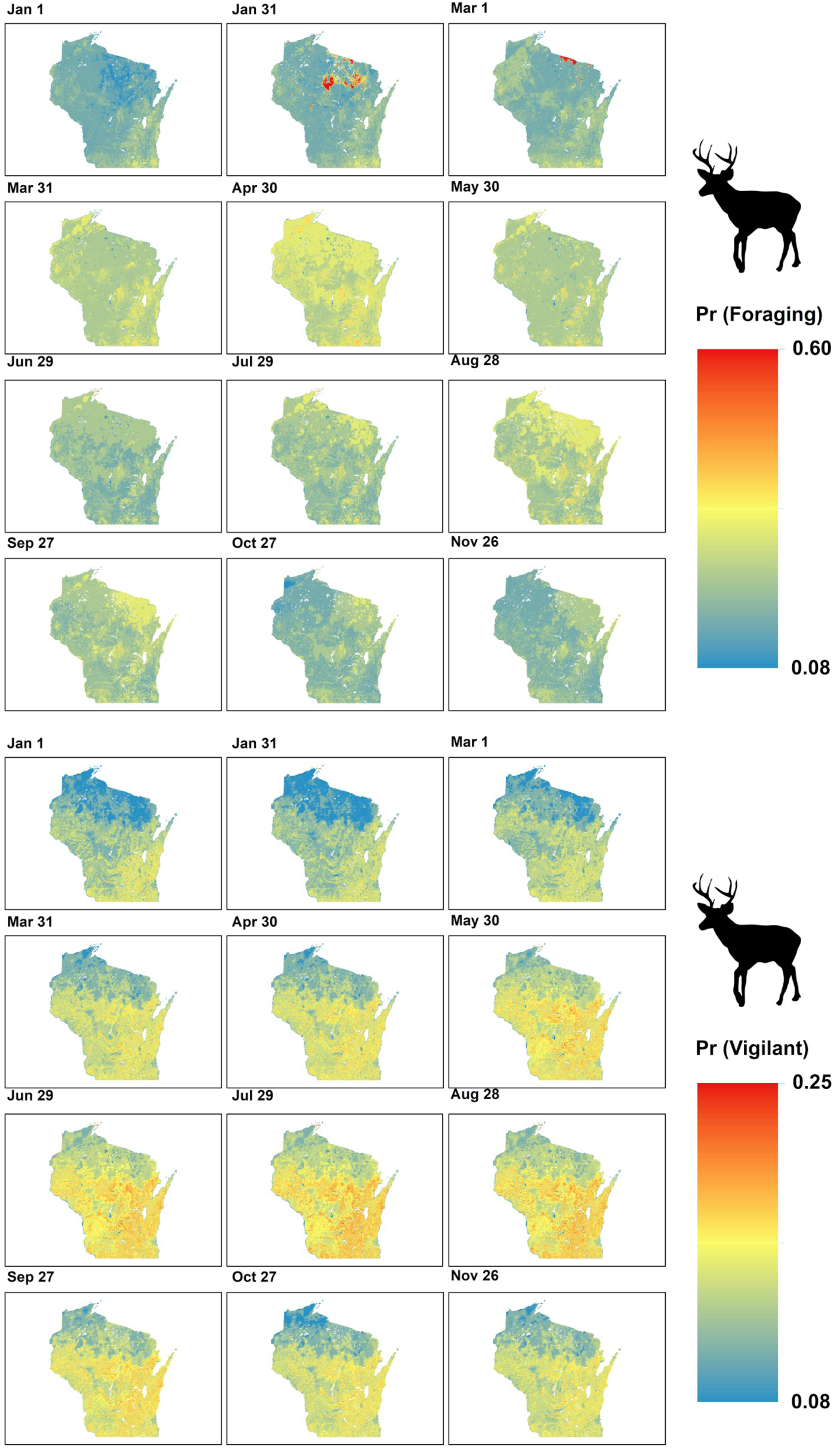
The predicted relative likelihood of deer foraging and vigilance behaviors (conditional upon occurrence and being detected) varies substantially across space and over the course of 2017, with highly outsized behavioral responses (e.g., top, Jan 31 and Mar 1) associated with ephemeral but dramatic weather events. During the year, deer in Wisconsin’s low productivity but predator-rich northern and central forests appear to spend relatively more time foraging and less time vigilant.

### Case 5. Strengthening inference via additional lines of evidence

To date, research drawing upon the growing body of biodiversity observations has largely focused on exploring patterns and changes in species distributions or populations at broader extents or finer resolutions than have traditionally been studied (e.g., Fink et al. 2010). At jurisdictional scales, broad-scale data collected using JON often coexist with information from more targeted surveys or experiments. There is a growing appreciation for synergies between data sources, and the development of broader monitoring networks may not only be useful for addressing new questions or issues, but for strengthening the integration of data types across multiple scales (e.g., Stenglein et al. 2015).

A simple but meaningful synergy afforded by SW is its ability to generate independent information that can be compared with existing lines of evidence, which may either clarify decisions via corroboration, or weaken the support for any specific decision if there is lack of concordance (Cook et al. 2012). For example, white-tailed deer are intensively monitored by WDNR using a population reconstruction model (Roseberry and Woolf 1991), but this model is implemented at a coarse resolution that make it challenging to draw linkages between population changes and potential drivers other than harvest regulations. The reliability of the current methodology has been questioned (e.g., Millspaugh et al. 2009), and there is interest in developing independent monitoring metrics. Detection-based metrics can be reasonable indices of deer abundance (e.g., Parsons et al. 2017) but are not always reliable monitoring metrics (Sollmann et al. 2013, Broadley et al. 2019). We contrasted harvest-based population estimates of deer in 2018 used by the WDNR with indices derived from concurrent SW data combined with remote sensing (Appendix S6) aggregated to the same county. Results (*r* = 0.55, Figure 6) suggest that each method can delineate broad spatial variation in deer density, and similar correlative strength across different years suggests harvest and SW data may be complementary monitoring data streams. SW permits prediction at finer resolution (Figure 6C), although predictions are not necessarily more accurate: regions of discrepancy are better viewed as areas where focused monitoring or further evaluation of the underlying model assumptions is warranted. For example, predictions from SW suggest higher deer density in northwestern Wisconsin than harvest-based models, which may reflect sampling discrepancies or confounding variation in detection rates driven by movement differences (Sollmann et al. 2013, Broadley et al. 2019). Both growing season phenology and varied land cover metrics showed associations with indices of deer abundance from SW, suggesting that growing season phenology could play a critical role in forecasting population changes to help guide harvest regulations (Hurley et al. 2014).

**Figure 6.**
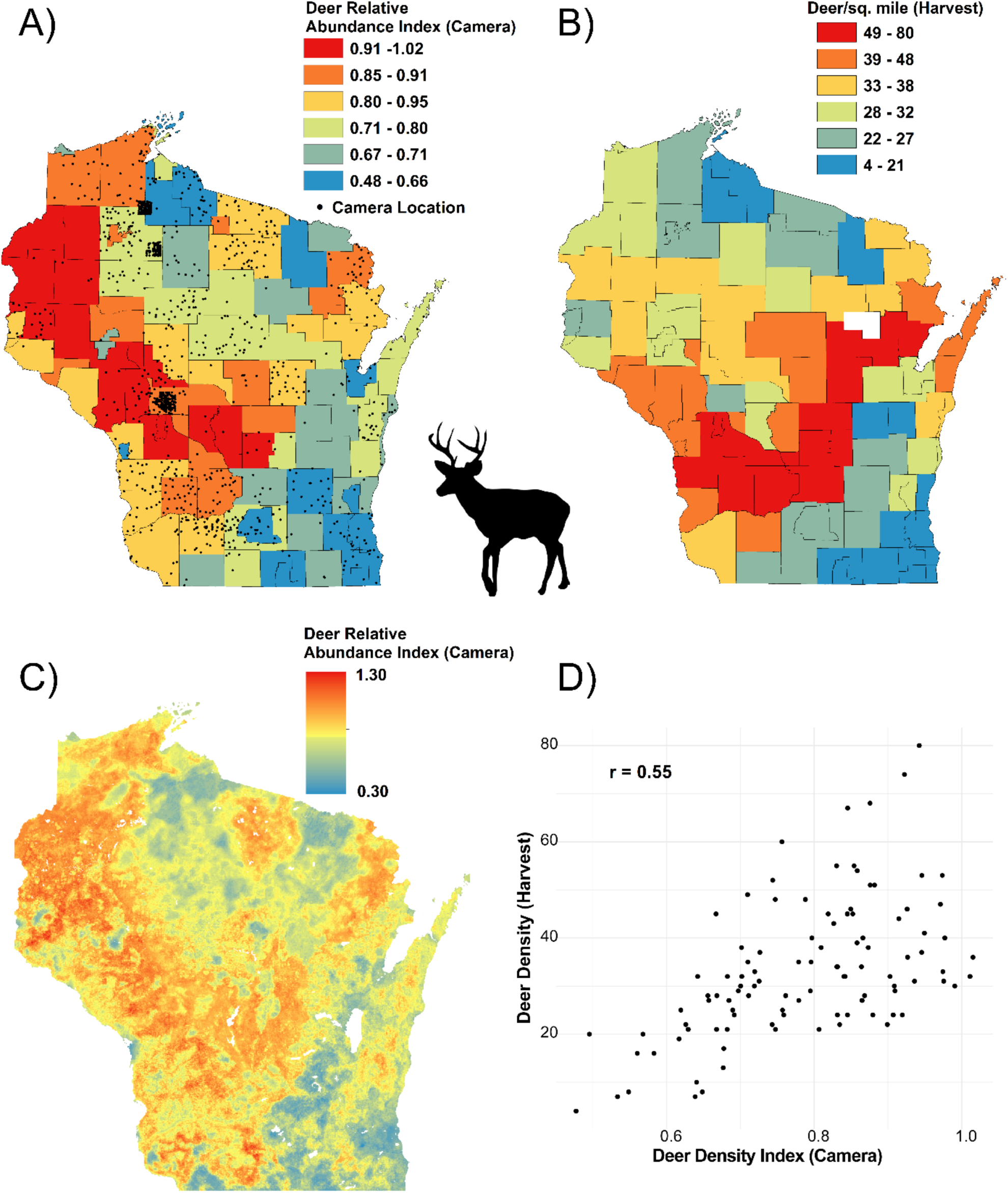
Patterns in 2018 deer relative abundance indices aggregated to the management unit derived from (A) prediction using Snapshot Wisconsin trail cameras and remote sensing, and (B) 2018 density estimates using a harvest-based population reconstruction model; (C) Spatially explicit (500 m cell) predictions used in (A); (D) Results from (A) and (B) are well correlated, although the potential resolution of inference or prediction is much greater using SW’s RAI (C) or more formal estimates.

Given the large number of species detected by SW, research is ongoing to assess SW’s role within WDNR’s existing monitoring efforts. In some cases, SW may complement or even replace existing monitoring efforts that are expensive, sensitive to survey conditions, or for which certain detection errors are more difficult to ascertain (e.g, aerial or snow-track surveys). Although empirical comparison of independent data streams can be a useful starting point, integrated models allow researchers to more formally identify or clarify specific methodological or analytical limitations and reconcile differences between approaches (e.g., Stenglein et al. 2015, Clare et al. 2017).

### Case 6: Scaling up inference using integrated modeling

Integrated models also provide an opportunity to ‘scale up’ information from local studies or estimate otherwise inscrutable parameters (e.g., Fletcher et al. 2016, Robinson et al. 2019, Sun et al. 2019). For example, bobcats are managed in two distinct zones in Wisconsin. The northern zone is monitored using an accounting model based upon harvested animals that has been subject to criticism (Allen et al. 2018b). The southern zone has been recently opened to harvest using results from a targeted population study (Clare et al. 2015), but the conservative quotas enacted to avoid overexploitation do not provide sufficient data for effective monitoring. Integrating detection/non-detection data from SW with previous bobcat capture-recapture data (Clare et al. 2015) enables estimation of population size and spatial variation in density. Results from an initial model reveal substantial spatial structure in bobcat density not possible using data from the earlier population study alone (Figure 7, methods in Appendix S7). Bobcats exhibit a pronounced range gap in southeastern Wisconsin that may reflect dispersal limitation and a slow recolonization process. Importantly, the model’s resolution allows population estimation within any spatial area of interest, and results suggest that the northern and southern zones have similar population sizes.

**Figure 7.**
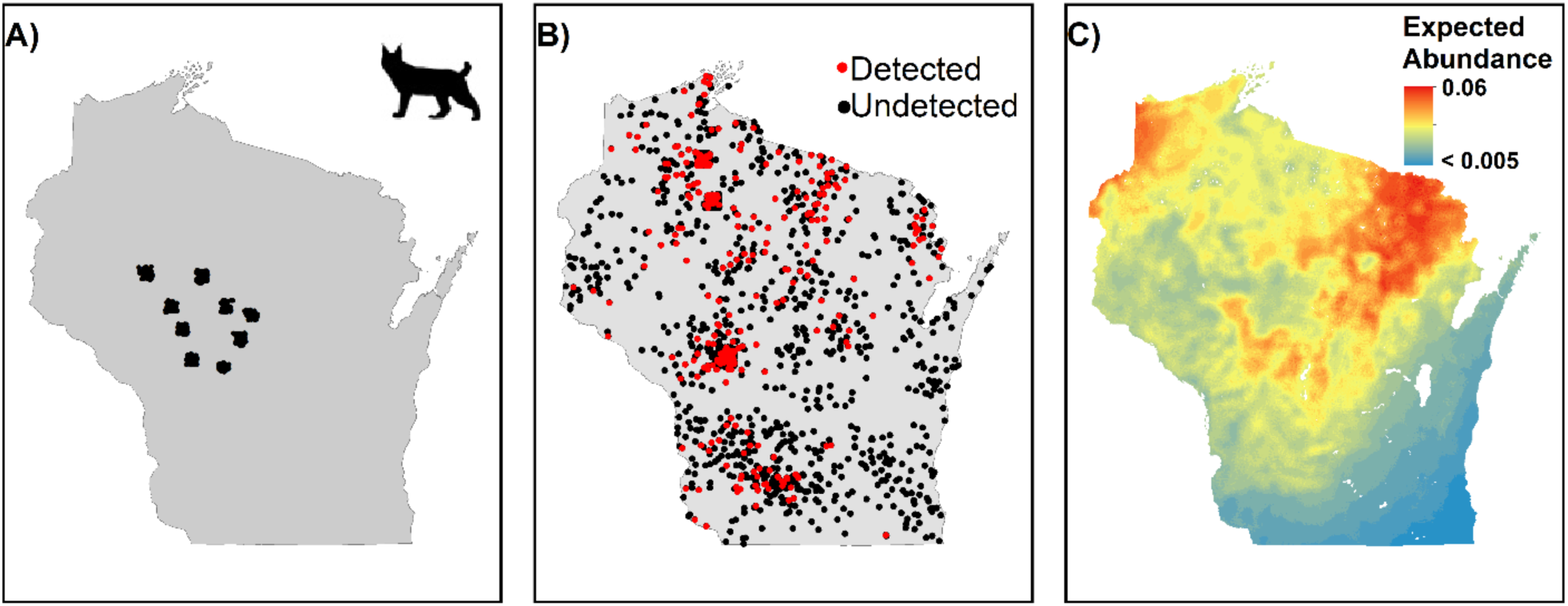
Smaller-scale bobcat studies (capture-recapture sampling, locations in (A)) were combined with the broader-scale SW sampling (locations in B) to scale up to state-wide bobcat density estimation (C).

## Discussion

Emerging technologies in remote sensing and ONs provide a powerful capacity to characterize biodiversity across both increasing spatial extents and finer levels of detail. The utility of SRS to planetary monitoring in an era of rapid anthropogenic change has been well established (Schlesinger 2006, Pettorelli et al. 2014, Jetz et al. 2016, Schimel et al. 2019). JONs operate using local forms of remote sensing data (such as citizen scientist observations, trail cameras, or passive acoustic monitoring) that complement SRS by quantifying variables that cannot be detected via traditional remote sensing platforms. These efforts facilitate discovery of new patterns and enable previously untenable lines of research (e.g., Kelling et al. 2009, La Sorte et al. 2018). SW represents the first example of an agency deploying a year-round observation network following a structured design across an entire state. Empowered by citizen scientists, SW has collected more wildlife observations during its short life span than all other monitoring programs focused on similar species in the state. SW demonstrates that the linkage of observation networks and SRS is tractable for agencies, and our case studies here demonstrate several ways in which monitoring data (and potentially management decisions) might be improved by actively pursuing such linkages.

The fusion of biodiversity observation and earth observation data through JONs can enhance performance across each primary role of science within the decision-making process: framing issues, delineating potential management actions, predicting the effects of these actions, and evaluating their efficacy (Keller 2009). JONs provide novel information about species, spaces, and times that is otherwise non-existent, which allows agencies to consider a broader set of emerging issues, and to consider and evaluate a wider range of management actions. JONs also provide information that can supplement or strengthen existing monitoring efforts by providing alternative lines of evidence, more rigorous or more precise estimation, and via integration with SRS, greater capacity to predict and understand variability in the state variables and functional relationships of interest. This enables agencies to more accurately predict the effects of potential actions and monitor the efficacy of management implementations with respect to issues of existing interest. Enhancing the dimensions of biodiversity observation also may provide agencies information needed to redefine overarching goals and objectives to more strongly align with spatial and temporal heterogeneity in stakeholder interests. Indeed, two of the most pointed challenges for wildlife management in North America involve generating a strong evidential basis for decisions and defining management goals and objectives that engage a broader range of potential stakeholders (Artelle et al. 2018).

The extent to which JONs can improve biodiversity monitoring and management depends on the degrees to which networks enhance the spatial, temporal, and biological extent or resolution over existing monitoring. The optimal sampling properties of a JON emerge from the spatiotemporal structure and dynamics of the organisms targeted and their environment, as well as properties of any existing monitoring. For example, Wisconsin’s seasonal environment leads to regular population cycles and relatively stable species distributions, and some processes of interest might be reasonably well-described with discrete annual dynamics. Other processes, such as those involving migratory, transient, or irruptive species in erratic or dynamic environments (e.g., Van Moorter et al. 2013), may require greater sampling extent and resolution to be effectively described.

Increased extent, resolution and integration of data will provide many serendipitous benefits. Many of SW’s potential contributions to research, monitoring, and management have yet to be realized. A few topics of particular interest include: a) delineating the distributions of specific physical traits and uncovering connections between population and trait dynamics (Zimova et al. 2020); b) fusing classified behavioral observations with the movement of tagged animals (e.g., Patterson et al. 2009) to enhance understanding of how the environment shapes space use; c) using biodiversity observations to estimate or fill spatial and temporal gaps in SRS (e.g., due to clouds) by leveraging estimated covariance between the two datastreams (Clark et al. 2017); and d) using biodiversity sensors as concurrent environmental sensors to evaluate or downscale spatial products derived from SRS (Siren et al. 2018, Hofmeester et al. in press). For example, SW cameras collect daily time-lapse images synched with the approximate overpass time of Landsat, Terra and Sentinel satellites to provide fine resolution observations of plant phenology and other environmental variables such as snow cover. Comparison with vegetation indices derived from SRS revealed that the trail camera phenology differed significantly (up to 17 days) depending on curve-fitting method and phenology metric, and that SRS indices did not consistently capture understory phenology (N. Liu, in prep.). Trail camera measurements may provide a systematic approach to disentangle understory and overstory from “land surface phenology” as measured using SRS. More broadly, the value of networks like SW may grow as technological and analytical capacity continues to improve.

Although local sensors can enhance the ecosystem measurements provided by SRS, in many other respects, locally-sensed data depend upon SRS’s spatially contiguous coverage for inference or prediction at meaningful scales. While smoothing methods can also be used for predicting across space or time, SRS provides contextual information that can potentially reveal details in patterns not necessarily apparent in raw observations. Some datasets like land cover products are commonly used by applied ecologists to predict or forecast varied state variables, but other products (e.g., the intensity of night-time lights or indices of plant productivity or phenology) are less widely used despite having potentially more direct connection to the variables of interest. On-going improvements in spatial and biological resolution of SRS data (i.e., broad-scale layers for vegetation structure and plant functional traits are near-term goals) will make it possible to generate new insights and better predict or forecast biodiversity responses. Similarly, as technology empowers increasingly continuous biodiversity sampling, both existing and nascent types of high temporal resolution environmental data provided by SRS may be necessary to explain spatiotemporal patterns in species distributions.

However, increased extent, resolution, and integration creates increased data volume within the observation network, which poses challenges. Informatics (data collection, classification, curation, computation) are limiting factors (La Sorte et al. 2018, Lindenmayer et al. 2018, Bayraktarov et al. 2019), as the size and complexity of the datasets generated pose challenges for data storage, management, and analysis. Technological solutions are improving rapidly for informatic concerns, including fast automated classification of images or audio recordings (e.g., Willi et al. 2019) and development of data portals that support curation (e.g., Sullivan et al. 2009) or provide classification services (Ahumada et al. 2020). In many cases, it will be substantially more efficient for programs to interface with existing cyberinfrastructure rather than create their own.

The novelty of the data produced by JONs poses additional challenges to articulating models for the systems of interest that can effectively inform management (Lindenmayer et al. 2018, Lindemayer and Likens 2018). A consequence of sampling and modeling species distributions at finer resolutions across greater extents is that the observations are driven by an increasing number of processes (local movement, density, lag effects associated with legacy processes) that operate at different scales (Yackulic and Ginsberg 2016). More predictors or new predictors, new process models, and new theories may be required to reasonably describe emergent patterns in the data (Coveney et al. 2016, Yackulic and Ginsberg 2016, La Sorte et al. 2018). In this regard, implementing JONs may require patience. Iterative prediction and testing processes may be necessary to refine theories and models fast enough to keep pace with the accumulation of data. JONs with more continuous sampling protocols may be best equipped to leverage the iterative forecasting cycle to improve understanding and management of the system of interest (Dietze et al. 2018). The evolution of eBird serves as a useful example. While earlier applications primarily relied on data-mining (Hochachka et al. 2008), the project has dedicated substantial effort towards gaining better understanding of the data collection process and mitigating potential observational errors (e.g., Hochachka et al. 2012, Johnston et al. 2018), developing models to exploit data dimensionality for improved synoptic ecological inference (Fink et al. 2010), and ensuring that the data could be employed for targeted applications (Robinson et al. 2018). As a project in an earlier stage of development, SW has itself dedicated substantial attention towards informatics and potential biases induced by data errors (Anhalt-Depies et al. 2019, Clare et al. 2019b, Locke et al. 2019), and our analyses here include some exploratory hypothesis generating techniques used when the phenomena in question was poorly understood. Where there was (is) sufficient previous research to guide a narrower set of hypothesis, we employed (and researchers should employ) analyses more suited to testing these hypotheses.

Finally, it is important to recognize that decision-making is a challenging process that requires balancing ecological information with other considerations. More or improved information does not necessarily lead to improved decision-making, as it can take time to determine how to weigh new findings, information may not exist at the time scales needed by decision-makers, or information may be poorly communicated (e.g., Sarewitz and Pielke 2007, Fuller et al. 2020). Similarly, factors other than ecological information may take precedence, and in other cases, the speed with which new ecological information becomes available or its volume can make it difficult for the decision-making apparatus to fully absorb it (e.g., McNie 2007). There has been increased recognition of the need to make products available more quickly to managers in a digestible form: for example, on-demand ‘real-time’ data access, processing, analysis and visualization are areas of substantial focus (e.g., Ahumada et al. 2020). However, agencies may lack the capacity or infrastructure to make decisions on-demand or in real time. Ultimately, reaching the full potential of JONs and other big data ecology may depend upon how quickly the managerial apparatus can evolve in its own right—for example, the capacity to more regularly make decisions and broaden the scope of management objectives.

Effective biological monitoring jointly quantifies changes in the system variables of interest, helps identify drivers of any changes in the system variables, and ultimately improves management (Nichols and Williams 2006, Lindenmayer and Likens 2010). Lack of data can limit conservation and management efforts, and the proliferation of data describing the earth’s surface and biodiversity upon the surface provides opportunities to vastly improve these efforts. Large scale deployment of remote sensors such as trail cameras provide great value by generating more information at increased spatial, temporal, and ecological resolution, and engaging citizen scientists to deploy sensors enables broadened sampling extents. However, even dense networks like SW are not spatially comprehensive; to be fully effective, JONs need remote sensing imagery and other geospatial data sets to maximize grain size and to permit effective hypothesis testing, prediction, and forecasting. JONs can further enhance both the taxonomic extent and the rigor of monitoring, enabling management agencies to break free from agenda driven or tautological monitoring (e.g., Clare et al. 2019a) and better achieve their missions.

## Acknowledgments

Support for this research was provided by NASA grants NNX16AO61H and NNX14AC36G, and a grant from the Federal Aid in Wildlife Restoration act awarded to Wisconsin Department of Natural Resources. We use data partially generated via the Zooniverse.org platform funded by a grant from the Alfred P. Sloan Foundation and a Global Impact Award from Google. Data and code used in this study are available at https://github.com/J-D-J-Clare/SWOverview.

## Supplemental information

Townsend, P.A., J. Clare, N. Liu, J.L. Stenglein, C. Anhalt-Depies, T.R. Van Dellen, N.A. Gilbert, A. Singh, K.J. Martin and B. Zuckerberg, submitted to *Ecological Applications* (2020). Integrating remote sensing and jurisdictional observation networks to improve the resolution of ecological management. bioRxiv supplemental materials.

## Appendix S1. Classification process details

Encrypted (initially unviewable) images from SW are uploaded by camera hosts to a central database using software developed internally at the Wisconsin Department of Natural Resources. After decryption within the central database, images can be viewed and classified by camera hosts on a secure agency hosted web-portal. Before images are made available for viewing and classification, a computer-vision based classifier attempts to screen out all images containing humans (agency personnel must check these screened images to ensure that wildlife images are not incidentally removed).

Camera hosts are expected to, at the least, tag any images of people, although they are encouraged to classify as many images as they like. Images that are not classified by camera hosts (or are classified by the host as something other than a limited set of species—Figure S1) are uploaded to the project’s Zooniverse hosted web platform (snapshotwisconsin.org). Visitors to the crowdsourcing platform view and classify image sequences (3 images associated with a triggering event) at random, and a specific image sequence is ‘retired’ from further classification once:

- Any three visitors classify the image as containing a deer
- If any 5 visitors agree on a species other than deer
- After 11 total classifications
- If one visitor has classified the image as containing a human
- If three visitors classify the image as being blank

For quality control, some images on the Zooniverse platform are classified by project professionals whom, when certain of the classification, can denote the image classification as ‘gold standard’. If available, the gold standard classification is treated as the final classification. Otherwise, the individual classifications are aggregated into a plurality consensus which is either treated as the ‘final’ classification or, for certain species, subject to a final expert classification. Note that non-expert ‘final’ classifications are stored in the database with a set of associated metadata previously shown to be good indicators of classification accuracy (Clare et al. 2019), so that data users can further screen out potential misclassifications as needed. The species specifically named in in Figure S1 are updated based upon evidence from more thorough but less regular reviews aimed at refining the estimates of classification accuracy. For example, recent work (C. Anhalt-Depies, unpublished data) has suggested that camera host classifications of striped skunks are sufficiently accurate (> 97%) that there is little benefit associated with subjecting these images to crowdsourcing, while gray foxes are inaccurately classified by both camera hosts and via crowdsourcing, and may generally require expert review. Similarly, while Clare et al. (2019) found that coyotes were relatively commonly misclassified via crowdsourcing, follow up work suggests that both camera hosts and crowdsourcing more accurately identify coyotes than previously recognized (C. Anhalt-Depies, unpublished data).

Although most of the analyses presented here and within the main text could have formally accounted for potential misclassification error (i.e., false positives--see Clare et al. in press), for simplicity, we either rely upon design-based solutions to reduce its incidence within the data used or ignore the error and accept that model outputs likely exhibit some bias.

---

**Figure S1.**
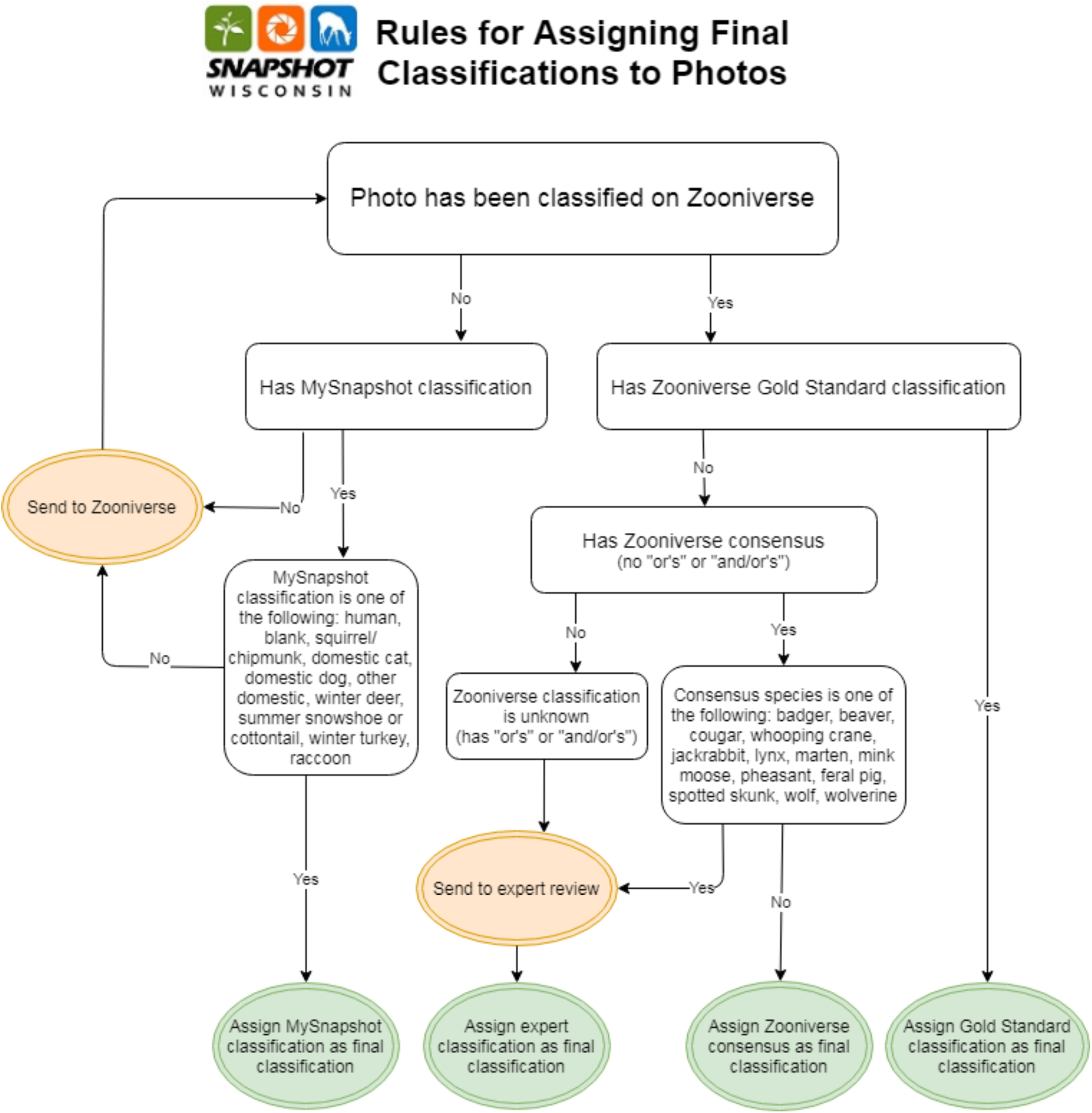
Image classification workflow employed by Snapshot Wisconsin.

## Appendix S2. Model Fitting Details for Bear Forecasting (Case 1)

To forecast changes in black bear distribution across Wisconsin and gain insight into the drivers of colonization and extinction processes, we fit a diffusion-based dynamic occupancy model (Broms et al 2016). Specific 5 x 5 km cells are indexed *i*, specific camera locations *k*, and specific years *t*. Succinctly, the model structure was as follows:

Detections *y*_*k,t*_ ∼ Binomial (Ndays_k,t_, *z*_*i[k],t*_ × *p*_*k,t*_) (Eq. S2.1), where Ndays_k,t_ is the number of 24 hour periods sampled by a specific camera in a given year, and where _*i[k]*_ indexes the cell a specific camera is located within. Some cells contained multiple camera locations: we assume here that cameras detect bears independently.

logit (*p*_k,t_) = α_0,k,t_ + α _1_Trail_k_ (Eq. S2.2), where Trail_k_ is a dummy variable denoting whether camera *k* was located along a maintained trail (e.g., a large hiking trail or forest road) or not.

logit (α_0,k,t_) ∼ Normal(μ_α_, σ_α_) (Eq. S2.3), i.e., the detection probability of each camera by year combination is subject to some extra logit-normal variation.

Latent occupancy state for a cell in a year *z*_*i,t*_ ∼ Bernoulli(*ψ*_i,t_) (Eq. S2.4).

logit (*ψ*_i,1_) = β_0_ + β_1_Forest5km_i_+ β_2_ForestWetland5km_i_ + *s*(*x, y*) (Eq. S2.5), where Forest5km_i_ and ForestWetland5km_i_ denote the proportion of these two land cover types within each cell (sourced from WiscLand 2.0), and *s*(*x, y*) is a cubic-spline smoother (with 30 basis functions) over X and Y coordinates (see knot locations in Figure S2.1).

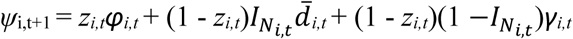 (Eq. S2.6), where *φ*_*i,t*_, 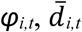, and *γ*_*i,t*_ denote the probability that an occupied cell remains occupied, that an unoccupied cell is locally colonized given that at least one neighboring cell was occupied in the previous time period, and the probability that an occupied cell is colonized from afar given that no neighboring cells were occupied in the previous time period.

logit (φ_i,t_) = δ_0_ + δ _1_Forest5km_i_+ δ _2_ForestWetland5km_i_ + δ_3_AC_i, t-1_ (Eq. S2.7), where AC_i, t-1_ is a centered autologistic term (see eq. 1 and description in Hepler et al. 2018) denoting the sum of the latent occupancy state across a neighborhood of cells in the previous time period (here, all cells within 12 km) minus the sum of the expected occupancy state across the same neighborhood in the previous year.

logit (γ_i,t_) = ω_0_ + ω_1_Forest5km_i_+ ω_2_ForestWetland5km_i_ (Eq. S2.8)

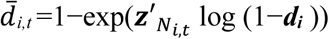 (Eq. S2.9), where 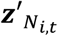 denotes the vector of occupancy states of cells neighboring *i*.

logit (***d***_***i***_) = θ_0_ + θ_1_**Forest5km**_**d**,i_+ θ_2_**ForestWetland5km**_**d**,i_ (Eq. S2.10), where the vector ***d***_***i***_ describes the diffusion gradient based upon a vector of pairwise differences in covariate values between cell *i* and its neighbors divided by the distance between a cell and its neighbors (e.g. **Forest5km**_**d**,i_); see eq. 8 - 10 in Broms et al. (2016). Here, we treat all neighbors (all cells within 12 km) as having equal distance, and do not divide covariate differences by distance.

Parameter estimates and uncertainty intervals are presented in Table S1. We fit the model using data from 1781 total camera locations across 889 specified 5 x 5 km cells between Julian days 150 and 270 in the years 2015 through 2018 (Figure S2.2). Total sampling effort was 232,300 24 hr camera sampling periods, with 4190 periods including a detection. We defined year 1 in the model as 2014, with years 2-5 informed by data sampled, and years 6 – 11 (i.e., 2019-2024) forecast using posterior simulation.

Increased sampling effort across years (Figure S2.2) appears to have made it challenging to estimate parameters associated with colonization, as none of the covariates used to model variation in γ_i,t_ or 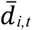 appeared to have any strong influence. Overall, local colonization appears to be slightly more likely than long distance colonization, and local colonization may lead to more stable growth because persistence appears to be more likely with occupied neighboring cells. Based upon previous work (Clare et al. 2019,

C. Anhalt-Depies unpublished data), the classification accuracy of black bears within images is believed to be very high (> 99% accuracy). Although the data used for fitting may include some false positives, we ignore such error here given previous evidence that its incidence is not likely to substantially skew model predictions and estimates. We did, however, review a small subset of classified images that appeared to be outliers based upon location and were expected to have more sizable leverage upon the results, and removed two clear false positives from locations in southeastern Wisconsin during 2017. Markov Chain Monte Carlo simulation was used to generate posterior samples across 4 chains using JAGS v. 4.0 (Plummer 2003). 10000 initial samples in each chain were discarded as burn-in, with 65000 subsequent samples per chain thinned by 50 to create a total number of 5200 posterior samples. (Code associated with varied case studies can be found at https://github.com/J-D-J-Clare/SWOverview).

---

**Figure S2.1.**
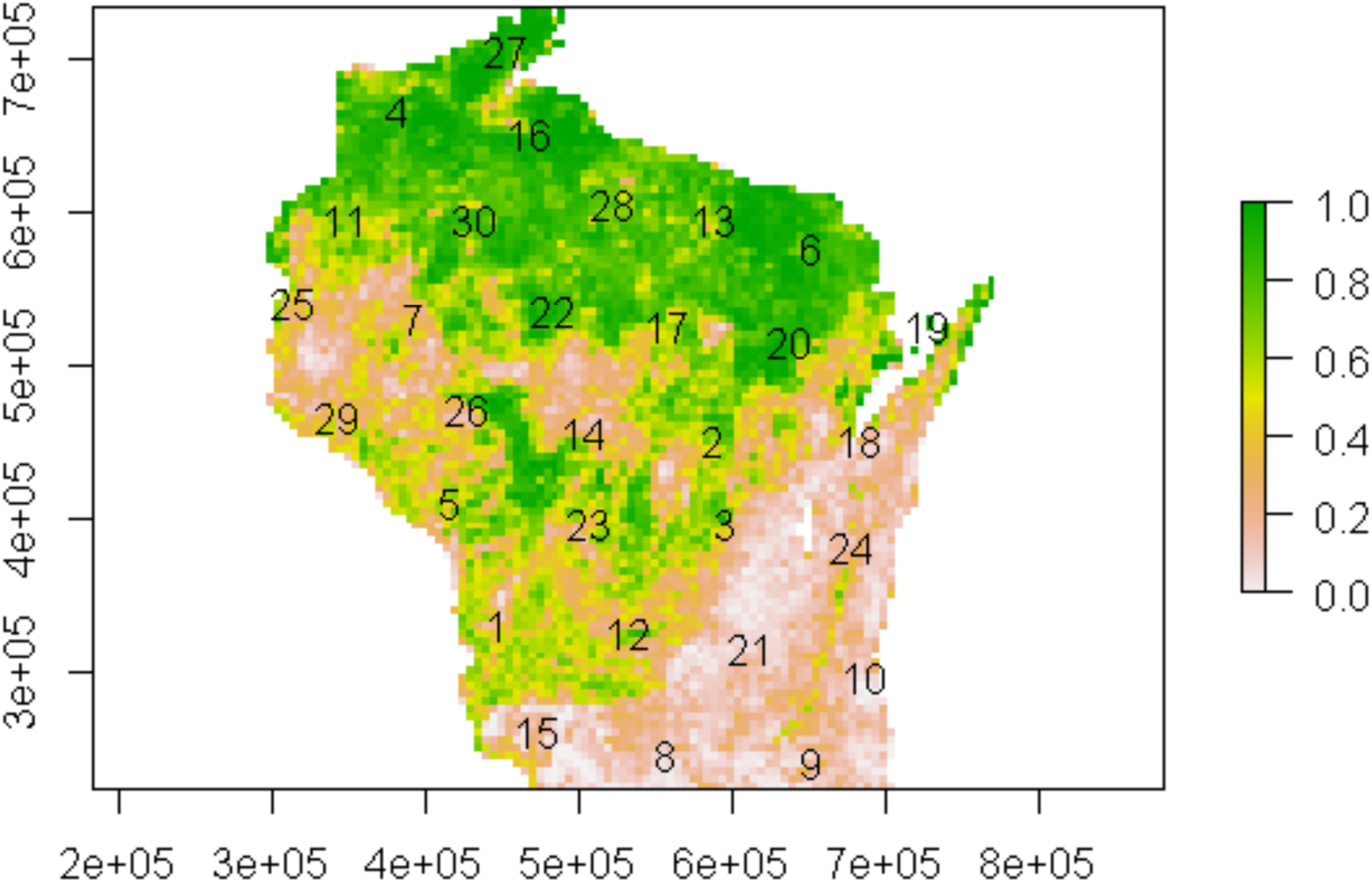
Proportion of forest land cover (forest + forested wetland) from Wiscland 2.0 across defined sites. Numbers indicate knot locations for spatial smoothing spline used to estimate initial bear occupancy.

**Figure S2.2.**
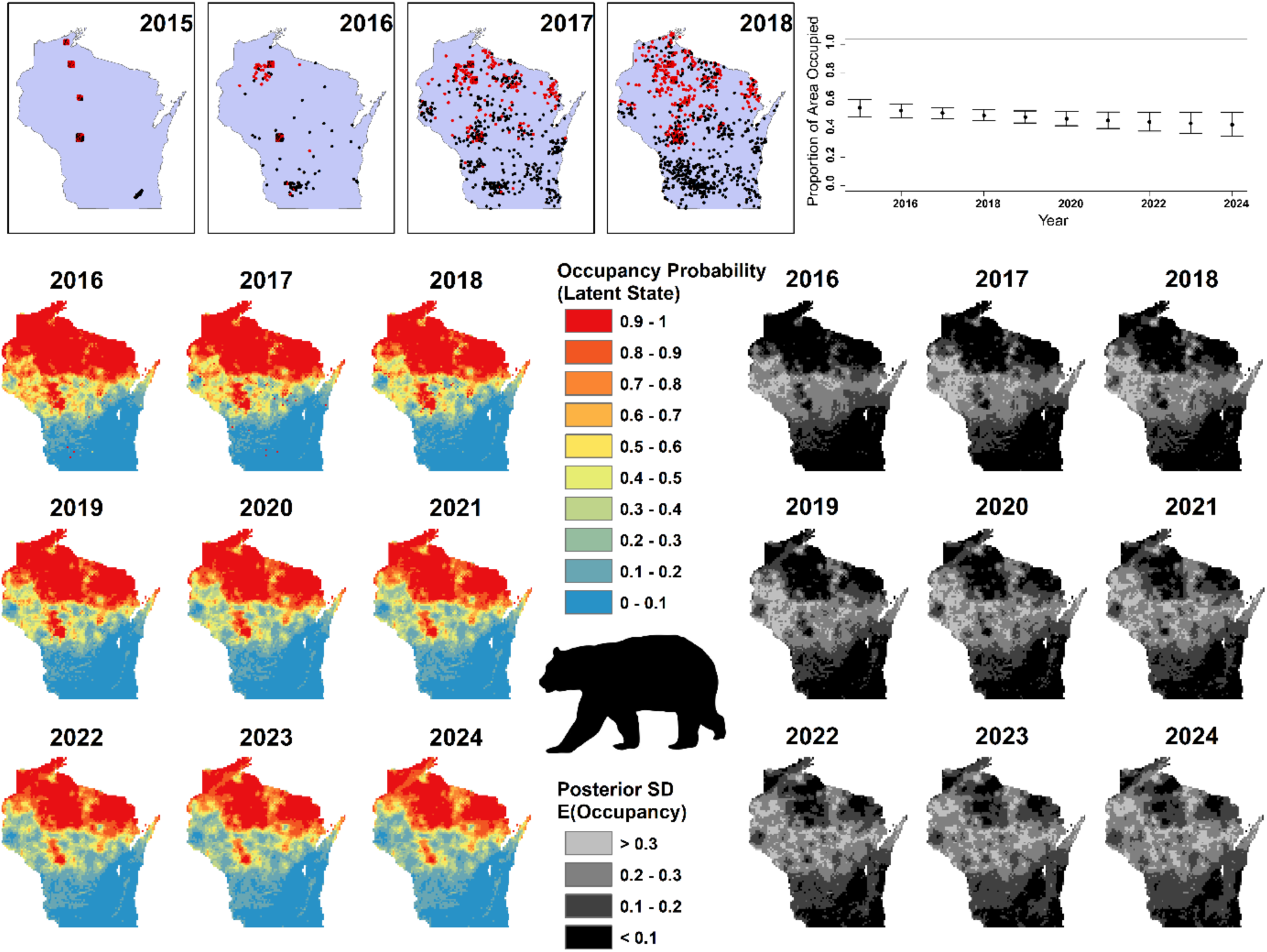
At top left, the spatial distribution of the sampling cameras, with red and black points denoting detection and non-detection, respectively. Top-right: estimated and predicted trend in the total proportion of area occupied by bears appears to be stable or perhaps slightly negative. Yearly estimates of the latent occupancy state (*z*) (bottom left) and the uncertainty in expected occupancy (*ψ*) (bottom right).

**Table S2.1.** Parameter estimates and uncertainty intervals associated with the diffusion-based dynamic occupancy model considered.

| Parameter | Posterior Mean | LCI | UCI |
| --- | --- | --- | --- |
| $\beta_0$ | 0.80 | -0.87 | 2.53 |
| $\beta_1$ | 0.99 | -0.07 | 2.11 |
| $\beta_2$ | 0.87 | -0.54 | 2.93 |
| smooth(x, y)<sub>1</sub> | -2.59 | -11.07 | 6.27 |
| smooth(x, y)<sub>2</sub> | -2.06 | -10.28 | 5.90 |
| smooth(x, y)<sub>3</sub> | -2.43 | -10.36 | 5.87 |
| smooth(x, y)<sub>4</sub> | 3.06 | -5.36 | 12.15 |
| smooth(x, y)<sub>5</sub> | -2.50 | -10.77 | 6.37 |
| smooth(x, y)<sub>6</sub> | 3.83 | -3.62 | 11.87 |
| smooth(x, y)<sub>7</sub> | 3.21 | -5.54 | 12.14 |
| smooth(x, y)<sub>8</sub> | -3.23 | -12.05 | 5.21 |
| smooth(x, y)<sub>9</sub> | -4.00 | -12.85 | 3.83 |
| smooth(x, y)<sub>10</sub> | -3.32 | -12.09 | 4.62 |
| smooth(x, y)<sub>11</sub> | 2.70 | -4.77 | 10.43 |
| smooth(x, y)<sub>12</sub> | -2.94 | -11.68 | 5.24 |
| smooth(x, y)<sub>13</sub> | 1.99 | -6.71 | 11.20 |
| smooth(x, y)<sub>14</sub> | 0.81 | -7.27 | 9.16 |
| smooth(x, y)<sub>15</sub> | -3.61 | -12.06 | 4.28 |
| smooth(x, y)<sub>16</sub> | 3.49 | -5.43 | 12.23 |
| smooth(x, y)<sub>17</sub> | 0.88 | -7.66 | 9.55 |
| smooth(x, y)<sub>18</sub> | -0.99 | -9.04 | 6.82 |
| smooth(x, y)<sub>19</sub> | 2.50 | -4.76 | 9.77 |
| smooth(x, y)<sub>20</sub> | 1.58 | -7.19 | 10.48 |
| smooth(x, y)<sub>21</sub> | -3.09 | -11.52 | 5.33 |
| smooth(x, y)<sub>22</sub> | 2.51 | -6.34 | 11.08 |
| smooth(x, y)<sub>23</sub> | -0.58 | -9.01 | 7.78 |
| smooth(x, y)<sub>24</sub> | -2.66 | -10.49 | 5.72 |
| smooth(x, y)<sub>25</sub> | 1.99 | -5.52 | 10.29 |
| smooth(x, y)<sub>26</sub> | 2.73 | -6.54 | 11.45 |
| smooth(x, y)<sub>27</sub> | 3.51 | -5.29 | 11.85 |
| smooth(x, y)<sub>28</sub> | 2.27 | -6.58 | 10.88 |
| smooth(x, y)<sub>29</sub> | 0.62 | -7.10 | 8.45 |
| smooth(x, y)<sub>30</sub> | 3.42 | -4.80 | 11.68 |
| $\theta_0$ | -3.85 | -5.07 | -2.92 |
| $\theta_1$ | 0.08 | -2.63 | 2.82 |
| $\theta_2$ | -0.59 | -3.62 | 2.43 |
| $\mu_\alpha$ | -4.52 | -4.62 | -4.42 |
| $\alpha_1$ | 0.68 | 0.50 | 0.87 |
| $\omega_0$ | -3.23 | -5.04 | -1.76 |
| $\omega_1$ | 0.60 | -0.51 | 1.75 |
| $\omega_2$ | -0.69 | -2.87 | 0.96 |
| $\delta_0$ | 3.93 | 2.73 | 5.23 |
| $\delta_1$ | 1.71 | 0.66 | 3.00 |
| $\delta_2$ | 1.71 | 0.45 | 3.10 |
| $\delta_3$ | 2.98 | 1.65 | 4.60 |
| $\sigma_\alpha$ | 1.25 | 1.18 | 1.32 |

## Appendix S3. Intra-annual Variability in Bear Occurrence (Case 2)

We used a variation of the multi-scale occupancy model described by Nichols et al. (2008) to generate predictions of the daily relative probability of bear occurrence across Wisconsin. The complete data-likelihood for the model is presented below because it may be more interpretable for some readers, and is structured as follows:

Let *z*_*i*_ denote the latent occupancy state across 500 m cells *i; z*_*i*_ ∼ Bernoulli(*ψ*_*i*_).

Logit(*ψ*_*i*_) = β_0_ + β_1_Forest5km_i_ + *s*(*x, y*) (Eq. S3.1), where Forest5km_i_ denotes the proportion of forest land cover within a 5 km buffer of each cell’s centroid and *s*(*x, y*) is a cubic-spline smoother over X and Y coordinates with 30 basis functions (following Appendix S2, see Figure S2.1).

Let *a*_*i,t*_ denote whether an organism is available to be detected in cell *i* during time period *t* = Julian Day 1, 2, 3, …365; *a*_*i,t*_ ∼ Bernoulli(*θ*_*i,t*_) (Eq. S3.2).

Logit (*θ*_*i,t*_) = *α*_*0,t*_ *+ α*_*1,t*_ Forest_i_+ *α*_*2,t*_ Cropland_i_ + *α*_*3*_EVI_i,t_ + *I*(t >1)*α*_*4*_*a*_*i,t-1*_ (Eq. S3.3); *α*_*0,t*,_ *α*_*1,t*_ and *α*_*2,t*_ were all derived as cyclical cubic splines with 15 basis functions. In other words, the model assumes a temporally varying intercept and temporally-varying logit-linear effects of the proportion of forest and cropland within cell *i*, a constant linear effect (*α*_*3*_) of MODIS enhanced vegetation index (EVI) in cell *i* during time *t*, and, for time periods beyond the first, a constant linear effect (*α*_*4*_) associated with availability in cell *i* during the previous time period.

Each camera (*k* = 1, 2, 3…*K*) was assumed to have a distinct probability of detecting a camera given that a bear was present and available for detection. We assume logit (*p*_*k*_) = δ_0,k_ + δ_1_Trail_k_ (Eq. S3.4), where δ_0,k_ ∼ Normal(μ_δ_, σ_δ_) (Eq. S3.5).

Observed detection/non-detection data at camera *k* on day *t*, denoted *y*_*k,t*_, was assumed to be distributed as Bernoulli(*z*_*i*[*k*]_ × *a*_*i*[*k*],*t*_ × *p*_*k*_).

Data came from 1,103 camera locations across 2017, and total sampling effort was 158,204 24-hr camera periods (where bears were either observed or not): predictions and maps in the main text are for the year 2017 as well. Without writing a custom MCMC sampler, existing software solutions using the CDL were prohibitively slow. However, the model above can also be formulated as a hidden Markov model, which can be fit far more quickly by marginalizing over the discrete latent states. Succinctly, we envisioned a 3 state HMM where s_i,t_ = 1 denotes that site *i* is unoccupied, s_i,t_ = 2 denotes *i* is occupied, but the species is not available during time *t*; and s_i,t_ = 3 denotes occupied and available for detection. We defined the initial state distribution as {[1− *ψ*_*i*_)] [*ψ*_*i*_(1− *θ*_*i,1*_)] [*ψ*_*i*_ *θ*_*i,1*_]}, where logit(*θ*_*i,1*_) = *α*_*0,1*_ *+ α*_*1,1*_ Forest_i_+ *α*_*2,1*_Cropland_i_ + *α*_*3*_EVI_i,1_ (Eq. S3.6).

Let **Γ**_*i,t-1*_ denote the state transition matrix used to define the state distribution at sites *i* for *t* = 2 through 365, where

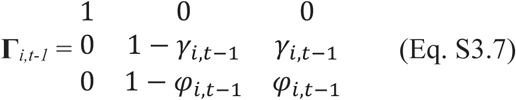

Above, logit(*γ*_*i,t*−1_)= *α*_*0,t*_ *+ α*_*1,t*_ Forest_i_+ *α*_*2,t*_ Cropland_i_ + *α*_*3*_EVI_i,t,_ (Eq. S3.8) and

logit(*φ*_*i,t*− 1_) = *α*_*0,t*_ *+ α*_*1,t*_ Forest_i_+ *α*_*2,t*_ Cropland_i_ + *α*_*3*_EVI_i,t_ + *α*_*4*_*a*_*i,t-1*_ (Eq. S3.9).

Let emission matrix **Ω**_**k**_ denote the probability of detecting the species (*y*_*k.t*_ = 2) or not (*y*_*k.t*_ = 1), where

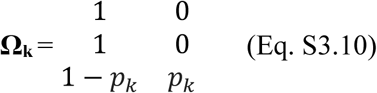

We the model by coding the forward algorithm and using Hamiltonian MCMC simulation through program Stan v2.18 (Carpenter et al. 2017, Stan Development Team 2018), with 2 chains of 1000 posterior samples used for inference after a burn-in of 1000 iterations. As with Case 1 (Appendix S2), we did not formally account for false positive error beyond evaluating a small set of observations that appeared to have been potential outliers, but we note that black bear classification has been estimated as being extremely accurate. Predictions in the main text reflect posterior prediction of pr(s_i,t_ = 3).

Results are presented in Table S3.1, with effects of interest plotted in Figure S3.1.

---

**Figure S3.1.**
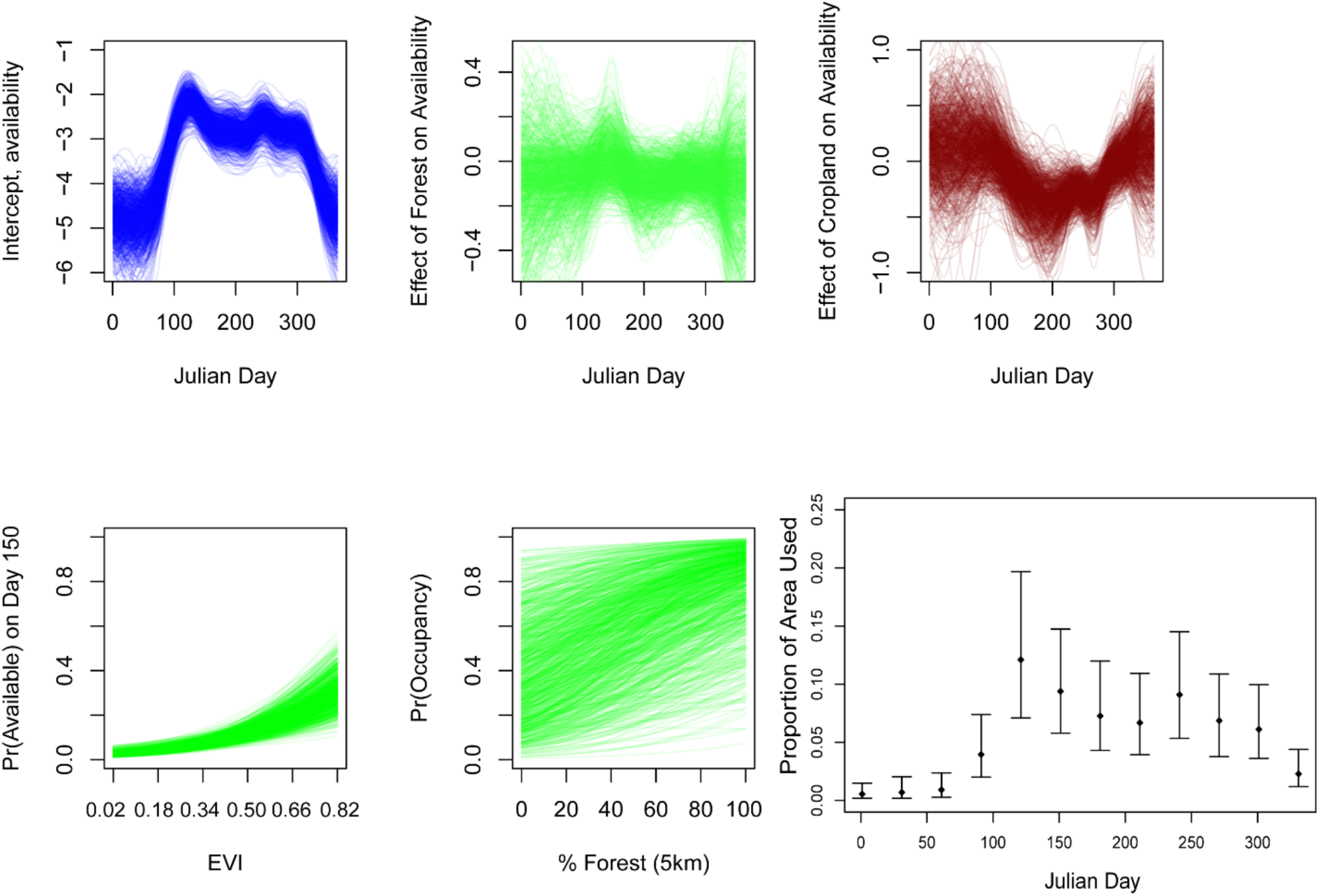
At top, temporally varying intercept (left) and effect of forest and cropland upon bear availability for detection (each line represents a posterior iterations). At bottom left and center, the effect of the MODIS Enhanced Vegetation Index (EVI) upon availability for detection (conditioning on day = 150 and all other covariates and mean values) and effect of % Forest upon cell occupancy. At bottom right, the proportion of cells across the state being ‘used’ on any given day exhibit strong cyclical dynamics consistent with bear phenology.

**Table S3.1.** Parameter estimates and uncertainty intervals associated with multi-scale occupancy model fit to predict intra-annual bear occurrence.

| Parameter | Estimate | LCI | UCI |
| --- | --- | --- | --- |
| $\beta_0$ | 0.58 | -1.59 | 2.75 |
| $\beta_1$ | 0.55 | 0.11 | 1.00 |
| s(X, Y)[1] | 1.29 | -1.71 | 4.40 |
| s(X, Y)[2] | -0.04 | -2.62 | 2.66 |
| s(X, Y)[3] | 1.24 | -1.48 | 4.15 |
| s(X, Y)[4] | -1.19 | -3.59 | 0.95 |
| s(X, Y)[5] | -0.70 | -3.41 | 1.82 |
| s(X, Y)[6] | 0.28 | -2.85 | 3.21 |
| s(X, Y)[7] | -0.07 | -3.03 | 2.71 |
| s(X, Y)[8] | 0.93 | -2.02 | 3.80 |
| s(X, Y)[9] | -0.68 | -3.04 | 1.46 |
| s(X, Y)[10] | -0.05 | -2.62 | 2.66 |
| s(X, Y)[11] | -0.60 | -3.46 | 2.22 |
| s(X, Y)[12] | -1.13 | -3.61 | 0.98 |
| s(X, Y)[13] | -1.34 | -4.16 | 1.24 |
| s(X, Y)[14] | -1.03 | -3.84 | 1.68 |
| s(X, Y)[15] | 0.45 | -2.44 | 3.25 |
| s(X, Y)[16] | 0.16 | -2.93 | 3.40 |
| s(X, Y)[17] | 1.19 | -1.52 | 3.91 |
| s(X, Y)[18] | 0.36 | -2.63 | 3.26 |
| s(X, Y)[19] | 1.29 | -1.76 | 4.27 |
| s(X, Y)[20] | -0.84 | -3.50 | 2.00 |
| s(X, Y)[21] | 1.07 | -1.89 | 3.93 |
| s(X, Y)[22] | 0.50 | -2.33 | 3.24 |
| s(X, Y)[23] | -0.59 | -3.49 | 2.32 |
| s(X, Y)[24] | 1.11 | -1.77 | 4.16 |
| s(X, Y)[25] | 0.81 | -1.56 | 3.14 |
| s(X, Y)[26] | -1.27 | -3.90 | 1.25 |
| s(X, Y)[27] | 0.80 | -2.33 | 3.84 |
| s(X, Y)[28] | 0.94 | -2.07 | 3.95 |
| s(X, Y)[29] | 0.79 | -2.36 | 3.85 |
| s(X, Y)[30] | 0.48 | -2.12 | 3.10 |
| $\sigma_\delta$ | 1.05 | 0.94 | 1.16 |
| $\delta_1$ | 0.53 | 0.27 | 0.84 |
| $\mu_\delta$ | -3.25 | -3.45 | -3.04 |
| s(t) intercept[1] | -4.40 | -5.24 | -3.54 |
| s(t) intercept[2] | -4.79 | -6.04 | -3.68 |
| s(t) intercept[3] | -4.83 | -6.06 | -3.75 |
| s(t) intercept[4] | -4.86 | -6.12 | -3.78 |
| s(t) intercept[5] | -4.46 | -5.56 | -3.59 |
| s(t) intercept[6] | -2.81 | -3.50 | -2.16 |
| s(t) intercept[7] | -1.92 | -2.56 | -1.33 |
| s(t) intercept[8] | -2.49 | -3.07 | -1.99 |
| s(t) intercept[9] | -2.88 | -3.48 | -2.29 |
| s(t) intercept[10] | -2.77 | -3.44 | -2.13 |
| s(t) intercept[11] | -3.03 | -3.60 | -2.46 |
| s(t) intercept[12] | -2.38 | -2.95 | -1.77 |
| s(t) intercept[13] | -2.87 | -3.58 | -2.32 |
| s(t) intercept[14] | -2.83 | -3.37 | -2.30 |
| s(t) intercept[15] | -2.90 | -3.59 | -2.28 |
| s(t) Forest[1] | -0.11 | -0.62 | 0.31 |
| s(t) Forest[2] | -0.11 | -0.59 | 0.29 |
| s(t) Forest[3] | -0.09 | -0.54 | 0.31 |
| s(t) Forest[4] | -0.08 | -0.45 | 0.26 |
| s(t) Forest[5] | -0.07 | -0.44 | 0.21 |
| s(t) Forest[6] | -0.07 | -0.38 | 0.16 |
| s(t) Forest[7] | -0.02 | -0.22 | 0.21 |
| s(t) Forest[8] | 0.02 | -0.18 | 0.34 |
| s(t) Forest[9] | -0.05 | -0.24 | 0.17 |
| s(t) Forest[10] | -0.11 | -0.36 | 0.08 |
| s(t) Forest[11] | -0.09 | -0.31 | 0.10 |
| s(t) Forest[12] | -0.08 | -0.28 | 0.09 |
| s(t) Forest[13] | -0.05 | -0.22 | 0.15 |
| s(t) Forest[14] | -0.07 | -0.26 | 0.12 |
| s(t) Forest[15] | -0.06 | -0.32 | 0.18 |
| s(t) Cropland[1] | 0.11 | -0.51 | 0.70 |
| s(t) Cropland[2] | 0.13 | -0.53 | 0.78 |
| s(t) Cropland[3] | 0.11 | -0.59 | 0.81 |
| s(t) Cropland[4] | 0.12 | -0.55 | 0.80 |
| s(t) Cropland[5] | 0.14 | -0.42 | 0.77 |
| s(t) Cropland[6] | 0.14 | -0.28 | 0.77 |
| s(t) Cropland[7] | -0.01 | -0.40 | 0.45 |
| s(t) Cropland[8] | -0.23 | -0.74 | 0.18 |
| s(t) Cropland[9] | -0.33 | -0.83 | 0.06 |
| s(t) Cropland[10] | -0.43 | -1.01 | -0.03 |
| s(t) Cropland[11] | -0.33 | -0.73 | 0.04 |
| s(t) Cropland[12] | -0.22 | -0.50 | 0.13 |
| s(t) Cropland[13] | -0.39 | -0.90 | -0.05 |
| s(t) Cropland[14] | -0.06 | -0.36 | 0.32 |
| s(t) Cropland[15] | 0.03 | -0.33 | 0.47 |
| $\alpha_3$ | 0.57 | 0.36 | 0.78 |
| $\alpha_4$ | 4.20 | 3.37 | 5.28 |

## Appendix S4. Distribution Models for Coyote, Striped Skunk and Opossum (Case 3)

We fit species distribution models for coyote, striped skunk, and opossum (and several other species) using boosted regression trees fit via the R library ‘dismo’ (Elith et al. 2008, Hijmans et al. 2017). Data used for fitting were drawn from 2,411 camera locations over a total of 69,1605 24-hr camera periods between 2015 and 2019. We ignore any temporal variability for this analysis, fitting models where the response was observed detection or non-detection (i.e., a Bernoulli distribution). We used the ‘gbm.step’ function to enact 10-fold cross validation to assist with model tuning (following recommendations by Elith et al. 2008).

The primary objective of this model fitting was to develop relatively accurate maps of the species’ distributions. For each species, there was relatively little pre-established research focusing on specific habitat associations. Moreover, for these species (and for other species of interest and other modeling exercises), the number of potential ‘static’ or ‘annual’ predictors easily derived from existing remote sensing products was fairly substantial, and we were further interested in trying to identify whether certain variables were consistently more supported than other (highly correlated) variables (e.g., NDVI vs. EVI, % Forest within a 1 km buffer vs. within a 5 km buffer, etc.). Given our existing priorities, we used 163 predictors: measures of effort (# trap nights), X and Y coordinates (Wisconsin Transverse Mercator), and 160 other environmental variables summarized derived from remote sensing and listed in Table S4.1.

**Table S4.1.** Static predictors derived from SRS used to fit gradient-boosted models for predicting Opossum, Coyote, and Striped Skunk distributions.

| Type | Source | Description | Metrics | Initial Spatial Resolution | Spatial Extent for Calculation | Aggregated Statistics | Temporal Resolution | Temporal Extent |
| --- | --- | --- | --- | --- | --- | --- | --- | --- |
| **Land Cover** | WiscLand 2.0 | Cover Type | % Type (21 types) | 30 m | 0.5 km pixel, 1, 5, 10 km radii | Mean | Static | 2015 |
|  | WiscLand 2.0 | Landscape Metrics | Class Richness (L3 classification) |  | 0.5 km pixel, 1, 5, 10 km radii |  |  |  |
|  |  |  | Shannon Diversity (classes at L3 classification) |  | 0.5 km pixel, 1, 5, 10 km radii |  |  |  |
|  |  |  | Aggregation Index |  | 0.5 km pixel, 1, 5, 10 km radii |  |  |  |
|  |  |  | Edge Density (Landscape, derived from |  | 0.5 km pixel, 1, 5, 10 km radii |  |  |  |
|  |  |  | L1 classes) |  |  |  |  |  |
| **Climate** | MOD10A2 .006 | Maximum Snow Extent (January) | Mean (%) | 500 m | 0.5 km pixel, 1, 5, 10 km radii | Mean | 8-day, used annual max (i.e., mean of annual max) | 2000-2017 |
|  | MOD11A2 .005 | Land Surface Temperature (January) | Mean | 1 km | 0.5 km pixel, 1, 5, 10 km radii | Mean | 1-day, used annual minimum (i.e., static product is mean of annual minimum( | 2000-2017 |
| **Land Surface Phenology \*** | MCD43A4 | NDVI, EVI | Start of growing season | 500 m | 0.5 km pixel, 1, 5, 10 km radii | Mean (Over 4 year period) | 8-day | 2014-2017 |
|  |  |  | End of growing season |  |  | Mean |  |  |
|  |  |  | Length of season |  |  | Mean |  |  |
|  |  |  | Minimum |  |  | Mean |  |  |
|  |  |  | Maximum |  |  | Mean |  |  |
|  |  | EVI | Annual Integrated |  |  | Mean |  |  |
| **Human Population** | DMSP-OLS | Nighttime Lights | Raw value | 500 m | 0.5 km pixel, 1, 5, 10 km radii |  | static | 2017 |

\*Described in detail below.

Most data sets used were higher level products distributed directly by NASA or other agencies. An exception was metrics of land surface phenology, wich we derived from NASA MODIS imagery. We downloaded available reflectance data (product MCD43A4) and calculated vegetation indices for each image. We subsequently fit the double-logistic model of Beck et al. (2006) to each annual time-series, and used the estimated parameters associated with the start and end of growing seasons and minimum and maximum values as inputs (as in Table S4.1 above). We used the model parameters to derive daily smoothed values of EVI, from which we calculated annual integrated metrics (for each year, the sum of all daily EVI values between the start of the growing season and the end of the growing season).

Model fit was adequate to strong based on cross-validation estimates of the ROC area under the curve for coyote (mean cross-validated AUC = 0.80, sd = 0.04), striped skunk (CV AUC = 0.76, sd = 0.03), and opossum (CV AUC = 0.90, sd= 0.02). The 10 most important variables for each species are summarized in Table S4.2—in each case (and for essentially every other species examined), the most important variable was sampling effort—although we do not interpret marginal variable effects or interactions here because inference using this methodology is tenuous, and our objective was not inference. The maps presented in the main text generally conform to expectations and reasonably delineate expected patterns in the relative distributions of coyote, striped skunk and opossum based on personal observation. While skunk and opossum are generally identified relatively accurately (e.g., 90-95% true positive rate) from trailcam photo classifications, coyotes pose challenges and are commonly misidentified (TPR ∼ 0.75). Predictions are in the main text are standardized as pr(species detected)|200 trap-nights of effort. As such, model predictions are essentially indices of species abundance and movement, with these rate-based estimates being less sensitive to false positives than estimates based in species occupancy (J. Clare unpublished data).

**Table S4.2.** The ten variables with greatest relative influence within gradient-boosted models for opossum, coyote, and striped skunk distributions.

| Species | Variable | Relative Influence |
| --- | --- | --- |
| Opossum | Sampling Duration | 17.28 |
|  | Latitude | 16.39 |
|  | % Coniferous (10 km) | 9.28 |
|  | % Cropland (10 km) | 3.26 |
|  | Mean January LST ( 10 km) | 1.93 |
|  | Mean January Max Snow Extent ( 10 km) | 1.68 |
|  | % Aspen/Birch (10 km) | 1.54 |
|  | Longitude | 1.46 |
|  | % Central Hardwoods ( 1 km) | 1.34 |
|  | % Cropland (1 km) | 1.21 |
| Coyote | Sampling Duration | 41.04 |
|  | %Developed (5 km) | 2.32 |
|  | %Wetland (5 km) | 1.76 |
|  | %Water (1 km) | 1.57 |
|  | SDI (Level 3, 500m) | 1.22 |
|  | %Herbaceous Wetland (5 km) | 1.17 |
|  | Mean Length of Growing Season, EVI (1 km) | 1.12 |
|  | % Grassland ( 10 km) | 1.09 |
|  | Mean Length of Growing Season, NDVI (1 km) | 1.08 |
|  | %Water (500 m) | 1.03 |
| Striped Skunk | Sampling Duration | 21.78 |
|  | Mean End of Growing Season, NDVI (1km) | 2.64 |
|  | % Grassland (500 m) | 1.96 |
|  | % Grassland (1 km) | 1.56 |
|  | % Grassland (5 km) | 1.44 |
|  | Mean Length of Growing Season, NDVI (5 km) | 1.42 |
|  | Aggregation Index, Landscape (5 km) | 1.4 |
|  | % Aspen/Birch (10 km) | 1.39 |
|  | % Wetland (5 km) | 1.29 |
|  | Mean Integrated EVI (1 km) | 1.27 |

## Appendix S5. Deer Behavior (Case 4)

Only images subject to crowdsourced classification from Zooniverse are tagged with behaviors. All images used for analysis were filtered such that we only used images in which one individual deer was in the image (to avoid modeling complications associated with multiple animals) and such that the predicted accuracy of the species classification was > 90% (following Clare et al. 2019; purpose being to avoid behavioral classifications based upon a limited number of votes). All images were drawn from 2017. In total, images used for the analysis (118,306) were drawn from 869 camera locations over 29323 24-hr camera periods; the analyses employed assume the number of images classified as foraging or vigilant within each period are binomial random variables. This implies the behavior in each image is independent. This may not hold for images in quick succession, but for the present analysis we ignore likely assumption violations, and accept that any predictions made probably under-report uncertainty.

A second known assumption violation relates to behavioral misclassification. We summarize the accuracy of foraging and vigilance behavioral classifications in Table S5.1 below based upon a review of 1200 classified images. Overall, agreement between the crowd-sourced classifications and post-hoc classifications was >90% for both categories. In principle, it is possible for an image sequence of a single animal to be tagged with multiple behaviors (e.g., an animal might exhibit vigilance in one image, and then commence moving). In our analysis, measures of agreement reflect classifications of single behaviors within a sequence (e.g., foraging or vigilant) rather than any combinations of classification behaviors. In some cases, the crowd vote was split; for the purposes of this analysis, we allocated ‘ties’ to ‘behavior not exhibited’. Again, we ignore the effects of misclassification here, being content with estimating and predicting ‘relative’ probabilities of foraging/vigilance.

**Table S5.1.** Confusion matrix of evaluation of crowdsourced behavioral classifications used to model and predict deer behavior conditional upon occurrence.

| **Vigilance** |  | Expert Vote |  |
| --- | --- | --- | --- |
|  |  | 0 | 1 |
| Crowd<br>Vote | 0 | 1049 | 44 |
|  | 0 or 1 | 9 | 14 |
|  | 1 | 7 | 103 |
| **Foraging** |  | Expert Vote |  |
|  |  | 0 | 1 |
| Crowd<br>Vote | 0 | 848 | 8 |
|  | 0 or 1 | 23 | 7 |
|  | 1 | 77 | 258 |

We used the ‘bam’ routine in R library ‘mgcv’ with discretized covariates (Wood et al. 2015, Wood et al. 2017, Wood 2017) to fit models for the probability that a trail camera image sequence was classified as ‘foraging’ or ‘vigilant’. Predictor variables were extracted from 500 m resolution pixels unless otherwise noted. Both models were fit with the same structure, which included:

- A multivariate tensor product smoother between latitude, longitude (each with 5 cubic regression basis functions), and day of the year (20 cyclical cubic basis functions), for the purpose of broadly capturing spatiotemporal structure.
- A camera-specific random effect (a smoother with a random effect basis, Wood 2017).
- Univariate smoothers with 20 cubic regression basis functions for the following variables:
  ∘ Integrated annual EVI, concurrent daily EVI (i.e., the EVI on the day of the observation), and concurrent daily EVI relative to neighboring pixels (i.e., the EVI within the pixel containing the camera on the day of the observation minus the mean EVI in the surrounding queen’s neighborhood on the same day). We viewed these variables as coarse proxies for vegetation productivity, and hypothesized that both foraging and vigilance behaviors might vary spatially as a function of annual productivity (i.e., integrated EVI), and might vary spatiotemporally depending upon the current state of vegetation productivity and its spatial juxtaposition.
  ∘ Landcover proportions (e.g., % cover) of Forest, Cropland, and Grassland+Hay vegetation classes (values derived from WiscLand 2.0).
  ∘ Landscape metrics: the edge density (across all level one classes), and the richness of level-three land cover classes. Both metrics calculated using WiscLand 2.0. Landcover richness was calculated within a 5km circular buffer. Edge and landcover richness capture finer grain heterogeneity in food resources (and presumably food phenology).
  ∘ The nighttime light intensity. We hypothesized that the intensity of nighttime lights—a proxy for proximity to human settlements -- would influence deer behavior with respect to human presence on landscape.
  ∘ Concurrent daily snow depth derived from SNODAS. Deep snow constrains deer movement, and we hypothesized that both foraging and vigilance in place would increase as a result.
- As a fixed effect, a dummy variable specifying whether the camera in question was placed along a maintained trail or not.

Estimated effects of the univariate smoothers are presented in Figures S5.1 and S5.2 below. Maintained trails had little effect on vigilance 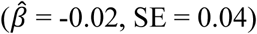, although deer appeared to forage slightly less frequently along maintained trails 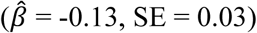. Overall, model fit for foraging (adj. r^2^ = 0.20, 18% of null deviance explained) was stronger than fit for vigilance (adj. r^2^ = 0.07, 7% of null deviance explained). For both models, the distribution of random site effects appears to have heavier tails than the assumed Gaussian distribution, and the proportion of variance explained by the predictors appears relatively limited compared to the random site effect or spatiotemporal terms. We expect that we are missing important potential predictors related to the occurrence or use-intensity of potential deer predators, and further expect that some of the covariate effects vary spatially or temporally. These topics, however, are the focus of ongoing research, and the results of this work are presented primarily as a demonstration of the integration of JONs and SRS data.

---

**Figure S5.1.**
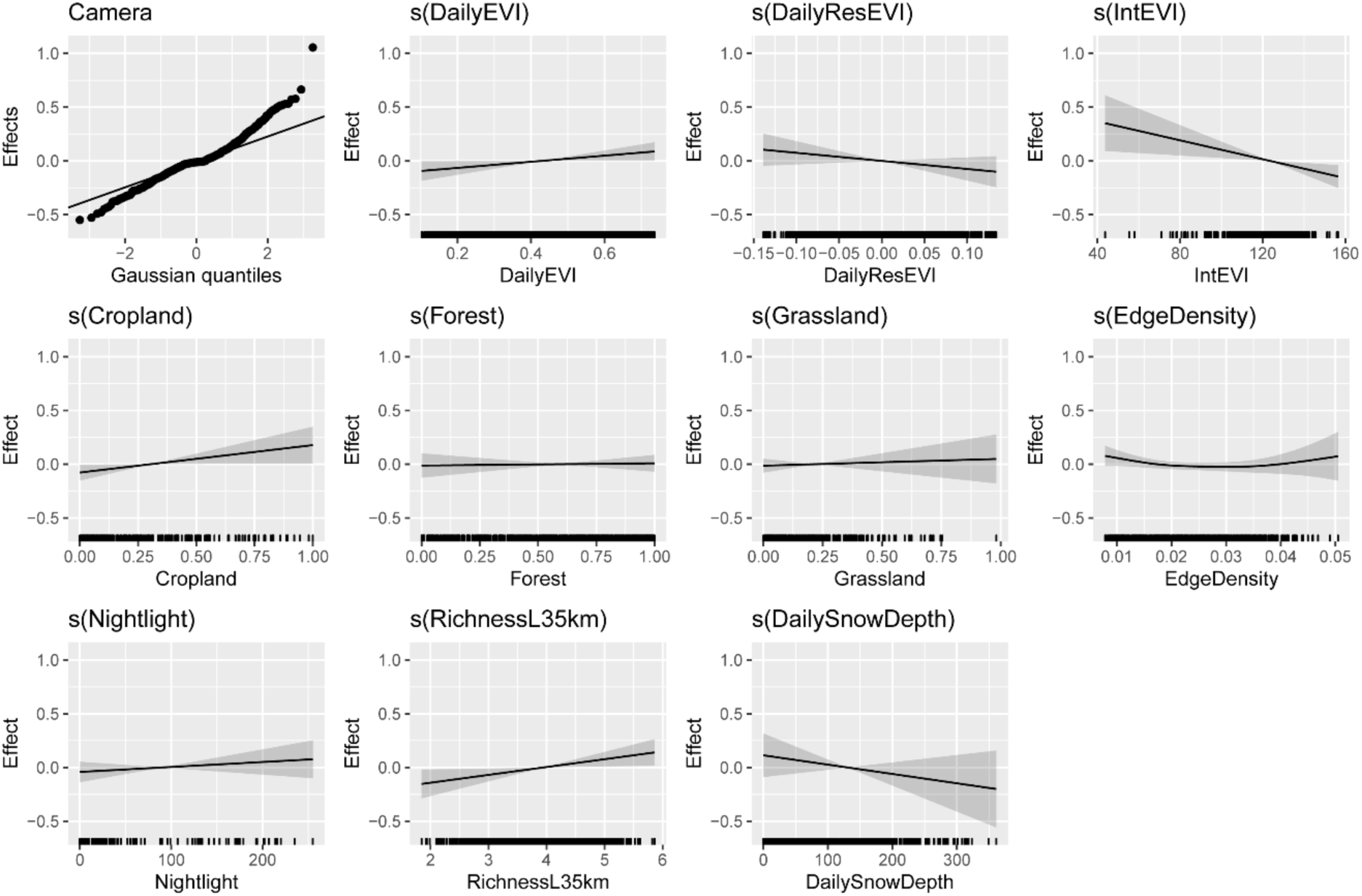
Effects of univariate smoothers on the probability of deer vigilance. Uncertainty intervals ignore uncertainty associated with the estimated intercept.

**Figure S5.2.**
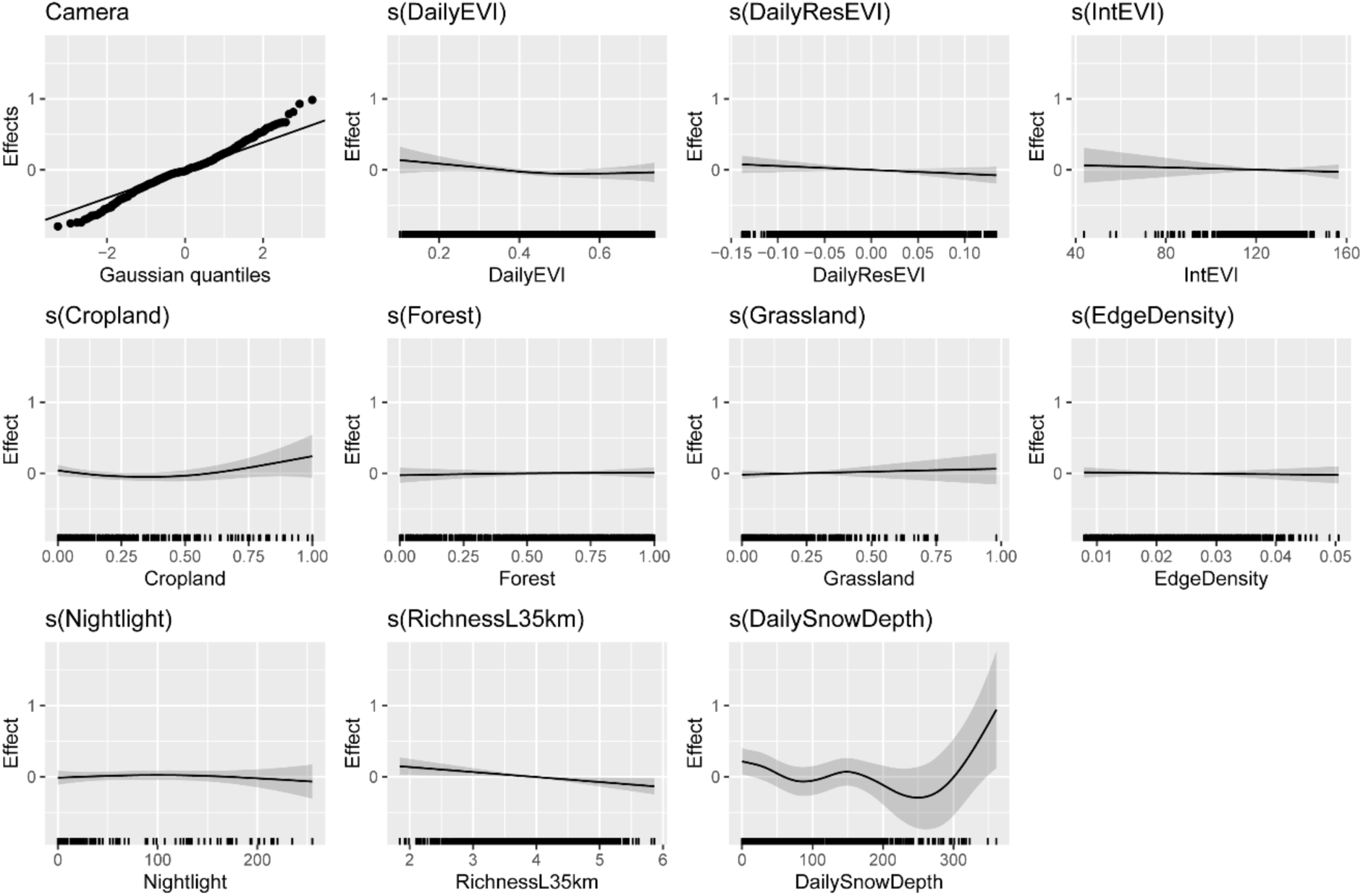
Effects of univariate smoothers on the probability of deer vigilance. Uncertainty intervals ignore uncertainty associated with the estimated intercept.

## Appendix S6. Comparison of Deer Relative Abundance Indices with Harvest-based Population Estimates (Case 5)

We used boosted regression trees (Elith et al. 2008) assuming normally-distributed residuals to generate predictions of the log catch per unit effort (e.g., log[detections]/days)—in the main text, we refer to this as a camera-based relative abundance index or a density index. The variables used for this analysis are the same as those presented in Table S4.1., with the ten most important variables for prediction of relative abundance presented in Table S6.1. The detection data used were drawn from 1105 camera locations across 98990 trap nights during summer (Julian Days 150 to 270) of 2018. Predictions (at 500 m resolution) reflect an exponentiation of the fitted value (which is on the log-scale).

Harvest-derived estimates used the sex-age-kill model (Roseberry and Woolf 1991) across Wisconsin’s deer management units, with a spatial-smoothing term used to account for autocorrelation between units in close proximity. The main text presents the correlation between the harvest-based point estimate (at the unit level) and a unit-level aggregation of the predicted index (i.e., the mean of all predicted pixels within the management unit).

**Table S6.1.**
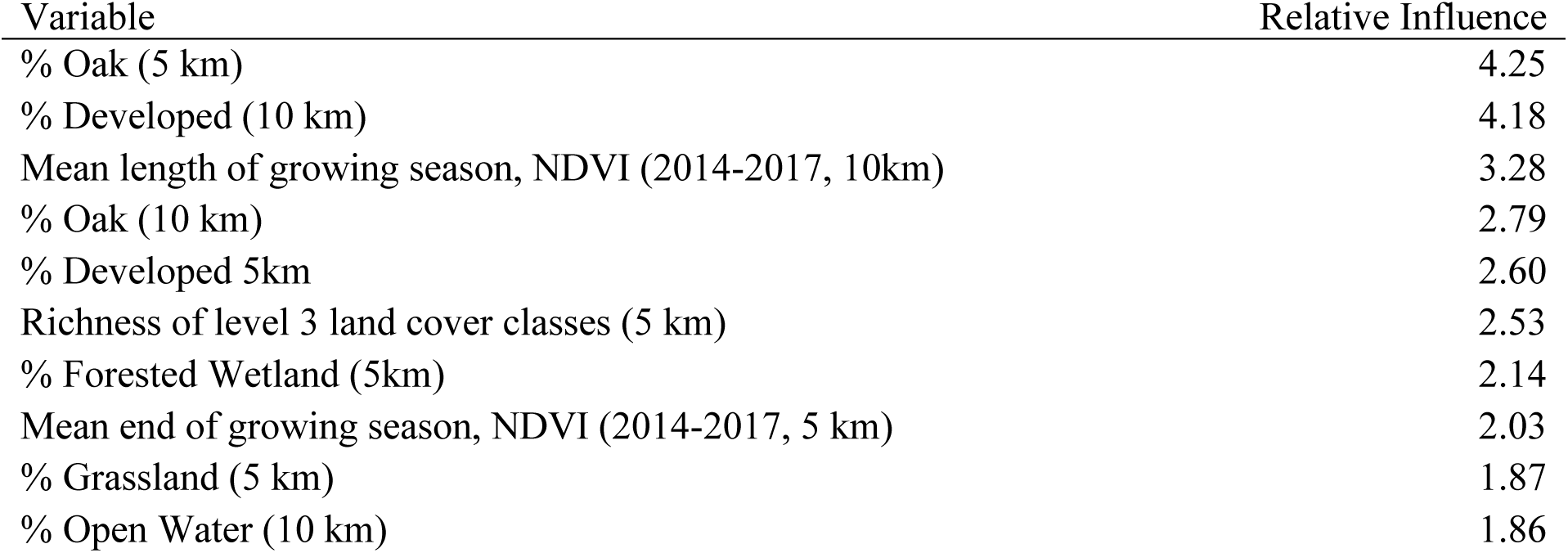
The (10) most important variables and influence scores used to generate predictions of white-tailed deer relative abundance across Wisconsin.

## Appendix S7. Estimating Bobcat population Size across Wisconsin (Case 6)

The analysis employed here was a spatially-explicit integrated population model (Chandler and Clark 2014) fit using a classical likelihood-based technique modified from Efford and Hunter (2018).

### Description of the model structure

The primary objective of the model is to estimate *µ*(***s*, β**), where *µ* denotes the intensity of an inhomogeneous point process for latent animal activity centers (see Borchers and Efford 2008, Royle et al. 2014), **s** denotes a vector of spatial locations (i.e., grid cells where an individual activity center might be located), and **β** denotes a vector of covariates used to model variation in *µ* across space. Spatial capture-recapture (SCR) models employ a detection function to describe a decay in the probability of observing an animal as a trap or detector is placed farther away from an individual’s activity center; here, we employ a hazard-half normal function where the expected count of detections for individual *i* at trap *j* on occasion *k* (*λ*_*i,j,k*_) is specified as:

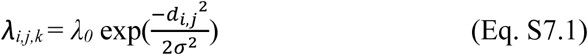

Here, *d*_*i,j*_ denotes the distance between the latent activity center of individual *i* and detector *j, λ*_*0*_ and *σ* are parameters to be estimated and may be themselves be functions of individual, trap, or time varying covariates (for example, log(*λ*_*0,t*_ = ***αX***). Throughout, we transform the hazard half normal function to describe the probability of >0 detections:

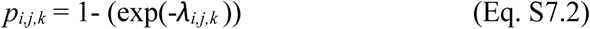

We assume that observed individual detection/non-detection at specific locations *y*_ijk_ is distributed as Bernoulli (*p*_*i,j,k*_).Let **θ** denote the full vector of detection parameters (*λ*_*0*_, *σ*, etc.).

Likelihood-based estimation of the detection and density parameters follows by maximizing the joint likelihood

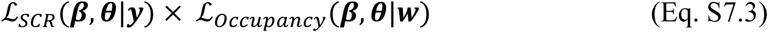

where ***w*** describes a vector of detection-nondetection data across traps and occasions where individuals are not distinguished. Our treatment of ℒ_*SCR*_(***β, θ***|***y***) follows the Poisson likelihood form described by Borchers and Efford (2008) and by Royle et al. (2014) on page 192, and marginalizes over the latent activity centers. Drawing from Efford and Hunter (2018, eqs. 4 and 11), we formulateℒ_*occupancy*_(***β, θ***|***w***) by assuming the expected count of detections for a particular detector *j* and occasion *k* can be described as

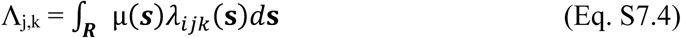

where ***R*** describes the gridded region of integration (i.e., the area assumed to contain the activity centers of all detected animals, identified or not). That is, the expected count of detections that a particular grid cell contributes towards a particular detector is equal to the product of the cell’s expected density and the expected number of times an individual with an activity center within the cell would be detected; the expected count of detections for any given detector is the sum of the expected contribution from each grid cell. The expected probability of > 0 detections at a specific location can be derived as

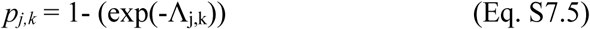

We assume *w*_*j,k*_ ∼ Bernoulli(*p*_*jk*_), or as we employ here, *w*_*j*_ ∼ Binomial (*p*_*jk*,_ *K*), such that

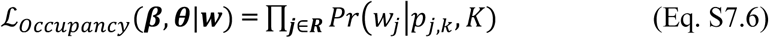

Note that Efford and Hunter (2018) focus on mark-resight estimation problems, and so their eq. 4 and 11 also include additional terms (e.g., to deal with the recruitment of unmarked individuals into a ‘marked’ population) that we do not need to consider here.

As noted by Efford and Hunter (2018), ***w*** is likely to be overdispersed. Strictly speaking, the generating process for a count of detections at a specific grid location is better described as ∫***R*** *n*(***s***)*λ*_*ijk*_(**s**)*d***s**, where n(**s**) describes the *realized* number of activity centers rather than the expected value. Because likelihood-based estimation must marginalize the latent activity centers out of the likelihood, the integral used for estimation uses the expected value, and observed Poisson or Binomial counts will be over-dispersed across trap locations relative to expectations.

Efford and Hunter (2018) propose maximizing a penalized Poisson or Binomial likelihood, with penalization informed by simulation. Our solution to this problem here is slightly different. First, we assume *w*_*j*_ ∼ Beta-Binomial (*g*_*jk*,_ *K*, δ), where δ is an overdispersion parameter to be estimated (Morris 1997). If the *J* detector locations contributing detection-nondetection data are more or less spatially independent (i.e., spaced far enough away from one another that the probability of a single individual being detected at two locations is negligible), simply estimating an over-dispersion parameter should be sufficient to properly estimate sampling variance without bias.

If the detector locations contributing occupancy data are not sufficiently spaced, the beta-binomial likelihood will not, on its own, be sufficient. In this case, we use a slightly different penalization for the occupancy data likelihood:

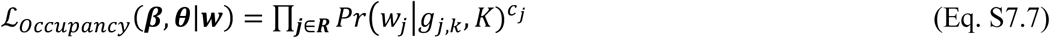

Here, we derive *c*_*j*_ as 1-*a* /*b*, where 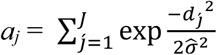 (Eq. S7.8), and 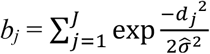 (Eq. S7.9), 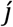 denotes all detectors other than *j*, and 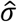 is an estimate of the σ parameter from a fitted model. The sum of *c* over all *j* detectors could be described as the effective number of independent detectors. For example, if there are only two traps (i.e., camera locations) producing occupancy data, and they are placed at the exact same location (but, say, at different times), the two ‘traps’ are collectively equal to one data point: ***a*** = (1, 1), ***b*** = (2, 2), and *c*_*1*_ and *c*_*2*_ both equal 0.5. More generally, the purpose of this penalization structure is to penalize traps with locational redundancy more heavily than traps placed in very distinct locations. Trap-specific effects upon sigma are fairly easy to accommodate (although not individual variation). Note that the occupancy data contribute very little information about σ (because the spatial structure of detection is discarded), and so penalization could be informed using estimates from a pure spatial capture-recapture (SCR) model.

This penalization, in many respects, is merely a very incremental step towards a more formal covariance structure for spatial patterns in the detection-nondetection data (Efford and Hunter 2018). These are straightforward to accommodate using Bayesian approaches that sample the latent *s*_*i*_ (Chandler and Royle 2013), but the data augmentation employed within such approaches can be prohibitively slow for large sampling problems unless highly customized samplers are employed.

In total, the fitting procedure is as follows. If the occupancy detectors are independent, maximize the joint likelihood ℒ_*SCR*_(***β, θ***|***y***) × ℒ_*occupancy*_(***β, θ***|***w***) using the Beta-binomial likelihood for ***w***. If the detectors are not spatially independent, maximize the pseudo-likelihood ℒ_*SCR*_(***β, θ***|***y***) × ℒ_*occupancy*_(***β, θ***|***w***)***c***. Note, however, that in many cases, the MLE appears to perform nearly as well as the MLPE.

### Simulation

We evaluated the estimators across 4 simulation scenarios depicted in Figure S7.1: each scenario employs the same landscape pattern with a single landscape covariate and a grid of 81 detectors spaced at 2 units producing SCR data in a fixed location over 20 sampling occasions. We fixed simulated abundance across the 100 by 100 unit lattice as 400 individuals, with the cell *g* containing each *s*_*i*_ simulated as Categorical (**π**), where 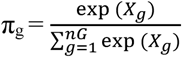 (Eq. S7.10). This implies covariate *X*_*g*_ has a log-linear effect of 1 upon density. SCR Detection parameters were simulated as *λ*_*0*_ = 0.135, and σ = 2. We assume the same parameters produce the detection/non-detection data—this does not need to be the case, although detection/non-detection data and SCR data must be able to assume either a shared detection process or a shared density process (Tourani et al. 2020). The general purpose of holding the landscape constant (and the location of the SCR detectors constant) is to avoid extra sampling variation associated with random landscapes (Efford and Fewster 2013) and potentially extremely sparse SCR datasets leading to unidentifiable parameters; we assume that SCR sampling aims to detect a reasonable number of individuals.

The placement of detectors (n = 81) collecting detection/non-detection data (across 20 occasions) varied by scenario, ranging from a fairly regular grid of detectors with trap spacing of 10 units (Figure S7.1 scenario A), to a tighter and less extensive grid with spacing of 3 units, to 9 clusters with intra-cluster spacing = 3 units (scenario C), and finally, to a much more comprehensive sampling grid (n = 441) with 3 units spacing (scenario D). Estimates and lognormal confidence intervals for expected population size and confidence intervals were constructed following Efford and Fewster (2013).

Both MLE and MLPE estimators described above were generally unbiased with respect to population size size, the covariate effect upon density, and detection parameters (Table S7.1); a slight negative bias with respect to total population size (that increases with effort) is consistent with simulation results presented by Efford and Hunter (2018). However, models fit using MLE--aside from case 1, in which the occupancy detectors were essentially independent—exhibited permissive coverage of both total population size and—particularly—the effect of the simulated covariate. The trap-varying penalized likelihood improved coverage, although it still fell slightly short of nominal in the simulation scenario D with the most intensive sampling.

**Figure S7.1.**
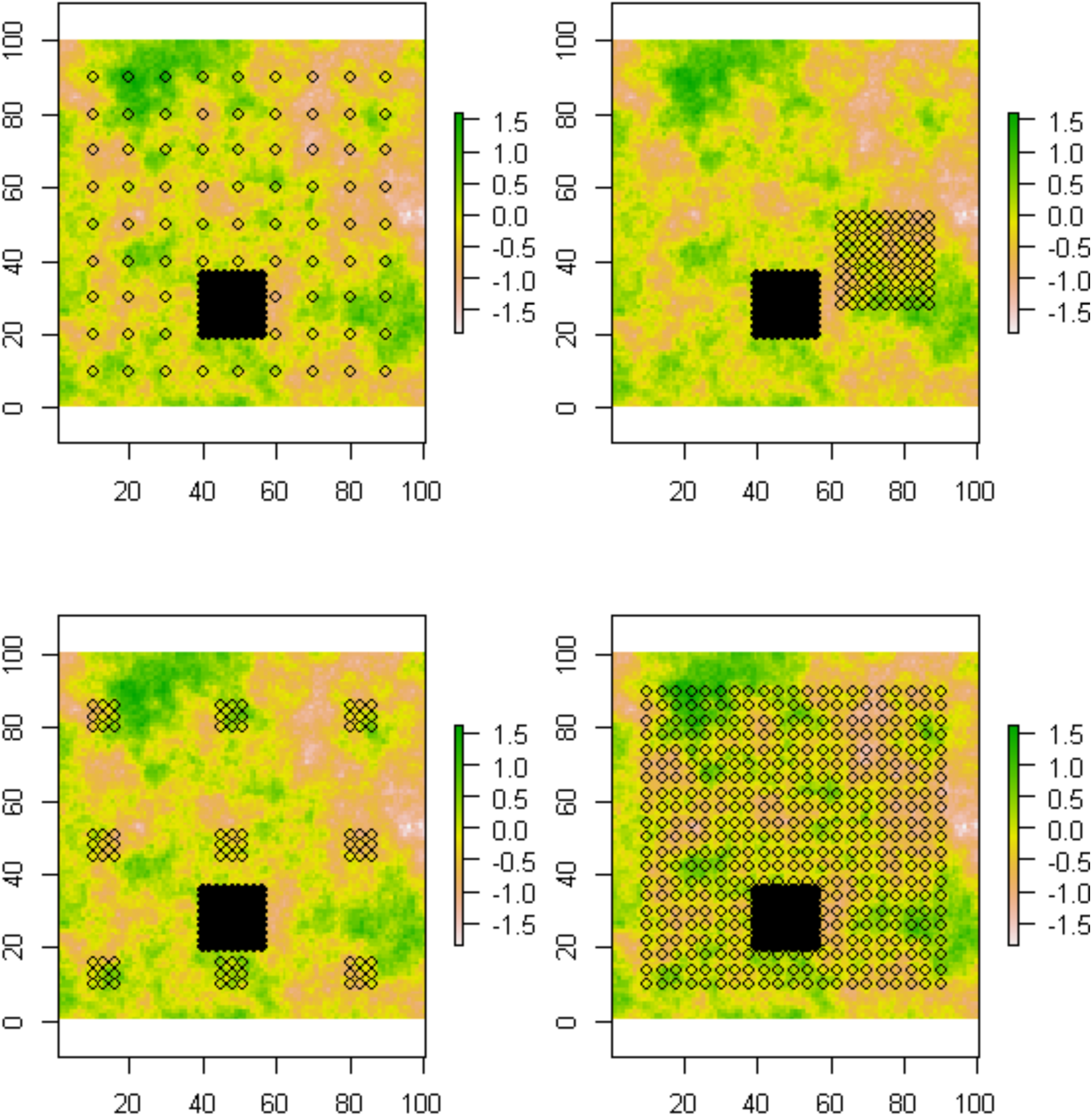
Sampling scenarios considered in the simulation study (clockwise from top left, Scenario A, B, D and C—SCR detectors are solid, occupancy type detectors are hollow. Coloration and scale at right denote covariate values associated with simulated landscape (greater values are associated with greater expected density). Map color indicates a single landscape pattern held constant for all scenarios.

**Table S7.1.** Simulation performance of maximum likelihood (MLE) and maximum pseudo-likelihood estimators (MPLE) with respect to total population size, the effect of a simulated covariates upon density, and detection parameters. RB = relative bias, RMSE = root mean squared error.

| Model | N | | | $\beta_1$ | | | $\lambda_0$ | | $\sigma$ | | # Traps | Effective<br># traps |
| --- | --- | --- | --- | --- | --- | --- | --- | --- | --- | --- | --- | --- |
|  | RB<br>(%) | Coverage | RMSE | RB<br>(%) | Coverage | RMSE | RB<br>(%) | Coverage | RB<br>(%) | Coverage |  |  |
| MLE A | -2.6 | 0.95 | 34.45 | -1.8 | 0.93 | 0.18 | 0.06 | 0.95 | 0.07 | 0.96 | 81 | (80.9) |
| MLE B | -1.3 | 0.87 | 44.02 | -0.3 | 0.766 | 0.33 | 0.04 | 0.95 | 0.1 | 0.96 | 81 |  |
| MLE C | -2.9 | 0.91 | 41.65 | -2.5 | 0.82 | 0.3 | 0.1 | 0.96 | 0.11 | 0.95 | 81 |  |
| MLE D | -3.2 | 0.92 | 28.12 | -0.81 | 0.85 | 0.12 | 0.1 | 0.94 | 0.27 | 0.97 | 441 |  |
| MPLE B | -1.2 | 0.95 | 41.06 | -1.5 | 0.93 | 0.3 | 0.06 | 0.96 | 0.09 | 0.96 | 81 | 32.4 |
| MPLE C | -2.3 | 0.98 | 40.22 | -2.9 | 0.94 | 0.29 | 0.1 | 0.96 | 0.09 | 0.96 | 81 | 39.83 |
| MPLE D | -3.1 | 0.93 | 28 | -0.86 | 0.92 | 0.12 | 0.1 | 0.94 | 0.24 | 0.97 | 441 | 279 |

### Case Study—Bobcat population estimation

Our case study integrates capture-recapture data in central Wisconsin collected in 2013 using trail cameras (Clare et al. 2015) with Snapshot Wisconsin data from 2018. That is, we make an explicit assumption that the expected density of bobcats across the area effectively sampled using SCR techniques and its spatial variability remained constant between these time periods. This assumption is likely not true in the strict sense, but we do not expect that bobcat density differed markedly between these time periods given that the species does not typically exhibit rapid population growth and that legal harvest of bobcats (within central Wisconsin) was extremely limited during this time period. Because integration required either assuming that detection or density parameters could be shared, and there was clear evidence that detection varied between the constituent SCR and SW surveys, we were willing to accept that the capability of generating a state-wide estimate came at the cost of some potential bias and loss of coverage. Data from SW were drawn from 1369 locations over 153,205 trapnights (between Julian Days 100 and 300): bobcats were detected on 809 of these occasions.

We modeled bobcat detection rates (λ0) as differing between trap locations along maintained vs unmaintained trails, and between the SCR and SW surveys given changes in equipment and protocols: log(λ0,j) = α_0,j_ + α_1_Trail_j_ (Eq. S7.11), where Trail_j_ is a dummy variables denoting whether a camera location was along a maintained trail, and where α_0,j_ varies depending upon whether a camera was associated with the SCR or SW survey effort. We assumed σ was constant.

We modeled variation in bobcat density as log(*µ*(***s***)) = β_0_+ β_1_Wooded_s_+ β_2_*sm*(Wooded_*S*_, σ_LS_)+ *s*(X_s_, Y_s_) (Eq. S7.12). Here, β_1_Wooded_s_ describes the focal effect of the proportion of a given cell s classified as a woody land cover type (Clare et al. 2015), and was derived from the 2011 National Land Cover Classification (NLCD, Homer et al. 2015). Term *sm*(X_s_, Y_s_) denotes a smoothing function over X and Y coordinates enacted via an unpenalized cubic regression spline with 10 basis functions (knots were selected using the cover.design function in R library ‘fields’; knots depicted in Figure S7.2). The term *sm*(Wooded_*S*_, σ_LS_) denotes a denotes a landscape smoother (Chandler and Heppinstall-Cymmerman 2016). The landscape smoother is essentially enacted by weighting values of the covariate Wooded_*S*_ across the spatial neighborhood of cell *s* (denoted *Z*_*s*_): *sm*(Wooded_*S*_, 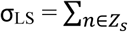 Wooded_*S*_*w*(*d*_*Sn*_, σ_LS_), where *w* denotes the weight of *s’* specific neighbor *n*, and where σ_LS_ is a parameter that controls the weight given the distance between *s* and *n* (*d*_*Sn*_). A half-normal based weighting scheme is derived as:

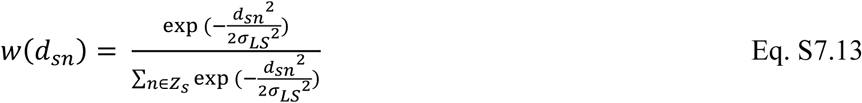

That is, coefficient β_2_ is applied to the weighted sum of the Wooded covariate across pixels within the neighborhood Z_s_ of cell *s*, ∑_*S*∈*Z*_ Wooded_*S*_*w*(*d*_*Sn*_, σ_LS_). Here, we defined Z_s_ for each pixel s as all other pixels in the region of integration *R* within 7 km (more broadly, we defined the region of integration used for fitting as the area within 7 km surrounding all camera locations; based on earlier fitting, this corresponds to roughly ∼ 4.5 times 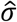). In fitting the model, we further normalized the weighted sum ∑_*S*∈*Z*_ Wooded_*S*_*w* (*d*_*Sn*_) by dividing it by the maximum value of the weighted sum across cells. The motivation for using the landscape smoother is both to estimate an appropriate extent at which the landscape covariate Wooded_*S*_ seems to have an effect (rather than estimate the effect within a predefined extent) and to account for the sensible hypothesis that landscape covariates have greater influence when closer to a focal site or animal activity center (Chandler and Heppinstall-Cymmerman 2016).

We fit the model using the optimizing function ‘nlm’ in R v 3.6. Parameter estimates and standard errors are reported in Table S7.2. Briefly, the results suggest that the baseline rate of encounter for SW surveys were less than for the SCR study and encounter rates were greater along maintained trails or logging roads. Estimates of σ and *σ*_*LS*_ imply that 50% and 95% of an individual’s detections are expected to fall within 1.80 and 3.75 km of the activity center, and that 50% and 95% of the overall neighborhood weights used to explain variation in density are estimated as occurring within 2.58 and 6.34 km from the focal cell. Note *σ*_*LS*_ was estimated with great uncertainty. The focal effect of woody cover was essentially negligible, while the amount of woody cover within the broader neighborhood was estimated as having a positive but extremely uncertain effect. The smoothing terms were also estimated with uncertainty, and imposing additional regularization (and adding knots) are ongoing focal points. Estimates of overall population size 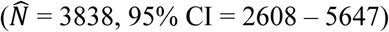 using the technique are lower than estimates derived from the accounting model currently used. Although it is unclear that the accounting model currently used is particularly accurate or reliable, the class of model we employ is known to exhibit negative bias if sources of variation in individual detection probability are not accounted for. In the near-term, evaluating alternative encounter models accommodating more detection heterogeneity is a similar goal. In sum, and the estimate from the model here is better viewed as a proof of concept, and is not intended to be used for decision-making.

---

**Table S7.2.** Parameter estimates and standard errors (SE) associated with empirical estimation of bobcat density across Wisconsin.

| Parameter | Estimate | SE |
| --- | --- | --- |
| $\alpha_{0,SCR}$ | -3.78 | 0.13 |
| $\alpha_{0,SW}$ | -4.66 | 0.24 |
| $\alpha_1$ | 0.80 | 0.11 |
| $\log(\sigma)$ | 0.43 | 0.06 |
| $\text{Log}(\sigma_{LS})$ | 1.04 | 2.37 |
| $\beta_0$ | -3.42 | 1.00 |
| $\beta_1$ | -0.06 | 0.44 |
| $\beta_2$ | 0.42 | 0.46 |
| $sm(X_s, Y_s) 1$ | 0.22 | 0.45 |
| $sm(X_s, Y_s) 2$ | 0.21 | 0.36 |
| $sm(X_s, Y_s) 3$ | 0.35 | 0.33 |
| $sm(X_s, Y_s) 4$ | -0.08 | 0.25 |
| $sm(X_s, Y_s) 5$ | -0.25 | 0.27 |
| $sm(X_s, Y_s) 6$ | -0.48 | 0.47 |
| $sm(X_s, Y_s) 7$ | -0.16 | 0.34 |
| $sm(X_s, Y_s) 8$ | 0.21 | 0.12 |
| $sm(X_s, Y_s) 9$ | -0.11 | 0.21 |
| $sm(X_s, Y_s) 10$ | -0.35 | 0.42 |
| $\delta$ | 3.82 | 0.14 |

**Figure S7.2.**
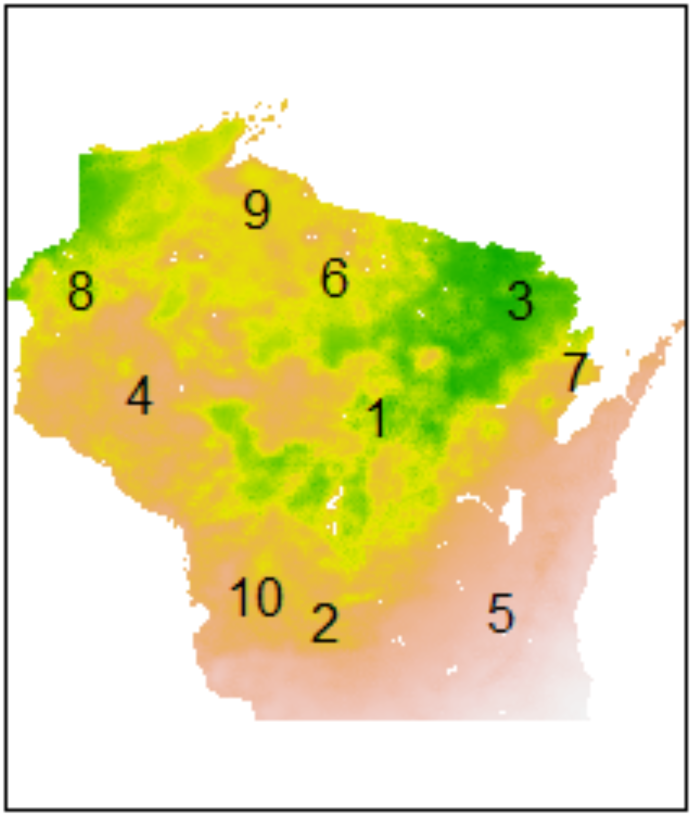
Location of knots used for spatial smoothing superimposed on gridded map of expected abundance prediction.

**Figure S7.3.**
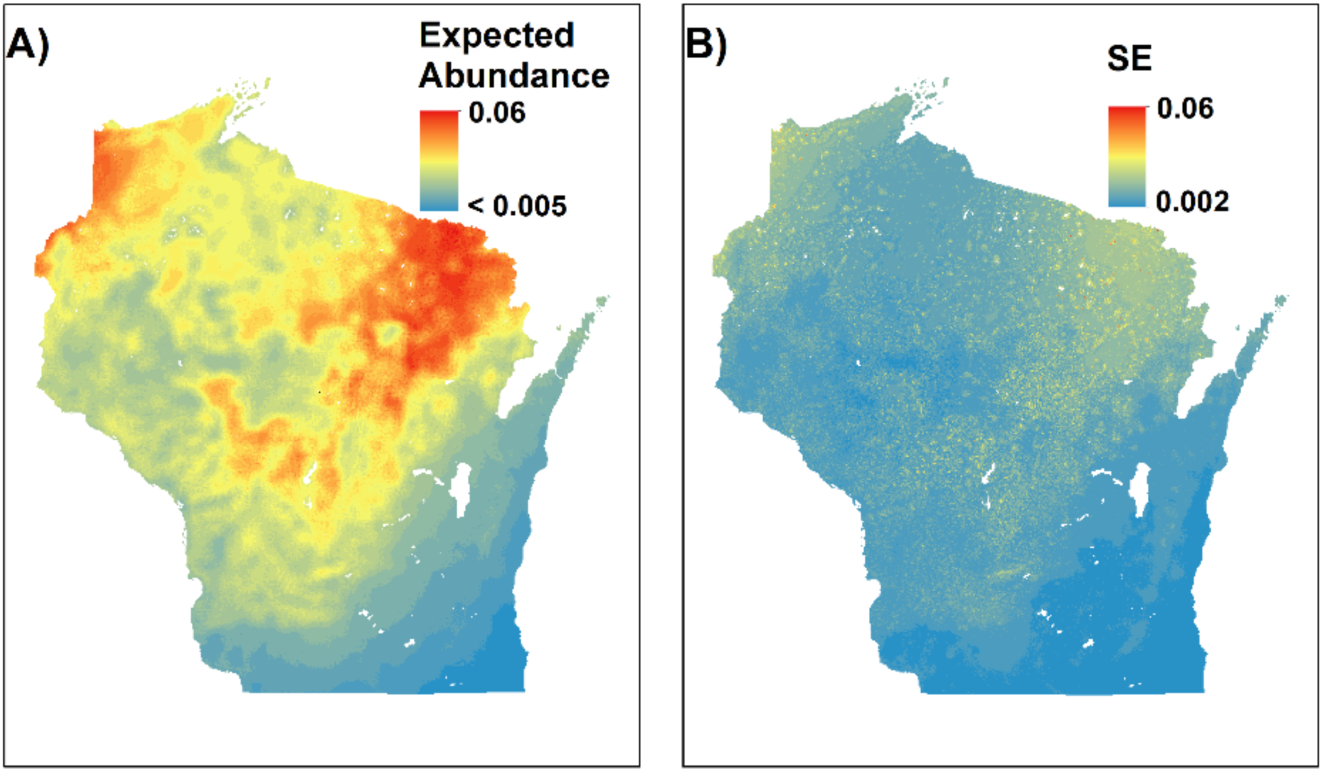
(A) Expected abundance (same as Figure 7 in main paper) and (B) standard error (uncertainty) from fitted model.

